# Cus2 enforces the first ATP-dependent step of splicing by binding to yeast SF3b1 through a UHM-ULM interaction

**DOI:** 10.1101/534347

**Authors:** Jason Talkish, Haller Igel, Oarteze Hunter, Steven W. Horner, Nazish N. Jeffery, Justin R. Leach, Jermaine L. Jenkins, Clara L. Kielkopf, Manuel Ares

## Abstract

Stable recognition of the intron branchpoint by the U2 snRNP to form the pre-spliceosome is the first ATP-dependent step of splicing. Genetic and biochemical data from yeast indicate that Cus2 aids U2 snRNA folding into the stem IIa conformation prior to pre-spliceosome formation. Cus2 must then be removed by an ATP-dependent function of Prp5 before assembly can progress. However, the location from which Cus2 is displaced and the nature of its binding to the U2 snRNP are unknown. Here, we show that Cus2 contains a conserved UHM (U2AF homology motif) that binds Hsh155, the yeast homolog of human SF3b1, through a conserved ULM (U2AF ligand motif). Mutations in either motif block binding and allow pre-spliceosome formation without ATP. A 2.0 Å resolution structure of the Hsh155 ULM in complex with the UHM of Tat-SF1, the human homolog of Cus2, and complementary binding assays show that the interaction is highly similar between yeast and humans. Furthermore, we show that Tat-SF1 can replace Cus2 function by enforcing ATP-dependence of pre-spliceosome formation in yeast extracts. Cus2 is removed before pre-spliceosome formation, and both Cus2 and its Hsh155 ULM binding site are absent from available cryo-EM structure models. However, our data are consistent with the apparent location of the disordered Hsh155 ULM between the U2 stem-loop IIa and the HEAT-repeats of Hsh155 that interact with Prp5. We propose a model in which Prp5 uses ATP to remove Cus2 from Hsh155 such that extended base pairing between U2 snRNA and the intron branchpoint can occur.

## INTRODUCTION

Pre-messenger RNA (pre-mRNA) splicing is a dynamic process catalyzed by the ribonucleoprotein complex known as the spliceosome. Assembly of the spliceosome, and subsequent catalysis and intron removal, involve numerous changing protein-protein, protein-RNA, and RNA-RNA interactions (Papasaikas and Valcarcel 2016; Fica and Nagai 2017; Scheres and Nagai 2017; Yan et al. 2019). Subunits of the spliceosome known as small nuclear ribonucleoproteins (snRNP)s, each named for the snRNA they carry, assemble in an ordered pathway of events. Many of these events are regulated in an ATP-dependent manner by DEXD/H proteins dedicated to specific steps in splicing (Chang et al. 2013; Liu and Cheng 2015). Upon ATP binding or hydrolysis, DExD/H proteins alter the binding of other components of the splicing machinery, promoting conformational changes within the RNA and protein components. These changes lead to release of some proteins and recruitment of others to the spliceosome, ensuring the fidelity and directionality of spliceosome assembly and catalysis (Koodathingal and Staley 2013). Recent cryo-electron microscopy (cryo-EM) models of spliceosomes at distinct steps (tri-snRNP, the A, B, B^act^, C, and P complexes) have brilliantly illuminated the composition and conformation of the spliceosome before and after each transition (Hang et al. 2015; Yan et al. 2015; Galej et al. 2016; Rauhut et al. 2016; Wan et al. 2016; Yan et al. 2016; Bai et al. 2017; Bertram et al. 2017a; Bertram et al. 2017b; Fica et al. 2017; Liu et al. 2017; Plaschka et al. 2017; Wan et al. 2017; Wilkinson et al. 2017; Yan et al. 2017; Zhang et al. 2017; Plaschka et al. 2018; Zhan et al. 2018b; Zhan et al. 2018a; Zhang et al. 2018). A challenge now is to understand the distinct mechanism of action of each DExD/H protein in catalyzing its specific transition. This challenge is more daunting given that several key spliceosomal components are part of early complexes that await structure determination, or are not present in the current models due to incomplete occupancy, transient association, or disordered nature.

The stable association of the U2 snRNP at the intron branchpoint (BP) to form the pre-spliceosome is the first event that requires ATP in splicing extracts from either yeast or mammalian cells (Will and Luhrmann 2011; Matera and Wang 2014). This event merges two large RNPs: the commitment complex (yeast) or the E-complex (human) containing the U1 snRNP bound to the 5’ splice site and a heterodimer composed of a pair of orthologous proteins (Msl5/SF1 and Mud2/U2AF) bound at the intron BP combines with the 17S U2 snRNP containing SF3a and SF3b proteins (Kramer 1996; Brow 2002). During this step, Msl5/SF1 and Mud2/U2AF are removed from the BP as it forms an extended duplex with U2 snRNA that bulges out a conserved adenosine for attack on the 5’ splice site. This duplex is packed into a crevice between Hsh155/SF3b1 and Rds3/PHF5A, where the bulged nucleotide occupies a pocket formed by both proteins in the A-complex (Plaschka et al. 2018). This ATP-requiring step appears to be mediated by the Prp5 protein, a DEAD-box family member with an RNA-stimulated ATPase activity important for its function in splicing (Ruby et al. 1993; O’Day et al. 1996; Abu Dayyeh et al. 2002; Perriman et al. 2003; Xu et al. 2004). One element of this ATP requirement is enforced by the nonessential U2 snRNP protein Cus2 (human homolog Tat-SF1) (Yan et al. 1998), because splicing extracts from Cus2 depleted cells can form pre-spliceosomes in the absence of ATP (Perriman and Ares 2000). Lethal ATP-binding site mutations of Prp5 are rescued by deletion of Cus2 (Perriman et al. 2003; Xu and Query 2007), suggesting that while it is not essential, removal of Cus2 from the U2 snRNP during A-complex assembly by an ATP-bound form of Prp5 is required for progression of spliceosome assembly.

Cus2 was initially identified as a suppressor of a cold sensitive misfolding mutant of U2 snRNA, and characterized as an RNA-binding protein associated with the U2 snRNP (Yan et al. 1998). During the spliceosome cycle, U2 snRNA conformation switches between two folded forms called stem IIa and stem IIc (Hilliker et al. 2007; Perriman and Ares 2007; Fica and Nagai 2017; Yan et al. 2019). To assemble into pre-spliceosomes, U2 snRNA must be folded into the stem IIa conformation (Zavanelli and Ares 1991; Zavanelli et al. 1994), but then is refolded to the stem IIc conformation prior to or during the first catalytic step (Galej et al. 2016; Rauhut et al. 2016; Wan et al. 2016; Yan et al. 2016). U2 snRNA dynamics are influenced by recombinant Cus2 *in vitro*, indicating that the protein can accelerate the interconversion between stem IIc and stem IIa (Rodgers et al. 2016). Together, these data suggest that Cus2 normally promotes recycling of the stem IIc form of U2 snRNA released after spliceosome disassembly to the assembly competent stem IIa form of U2 snRNA, but then must be released from the U2 snRNP by ATP-bound Prp5 for pre-spliceosome assembly. However, the location on the U2 snRNP where Cus2 binds, why its release from that location might be necessary for pre-spliceosome formation, and how the ATP-bound form of Prp5 promotes Cus2 removal are unknown.

Tat-SF1, the human homolog of Cus2, associates with the U2 snRNP (Fong and Zhou 2001; Agafonov et al. 2011) and has recently been shown to bind the U2 snRNP protein SF3b1 through a U2AF homology motif (UHM) in Tat-SF1 and a U2AF ligand motif (ULM) in SF3b1 (Loerch et al. 2018). UHMs adopt an RNA recognition motif (RRM)-like fold but have diverged in ways such that they no longer bind RNA, but rather interact with ULMs (Kielkopf et al. 2004; Loerch and Kielkopf 2016). These motifs have been shown to mediate binding between a number of early mammalian splicing factors including the BP-binding protein SF1, the polypyrimidine tract-binding heterodimer U2AF, and a number of alternative splicing factors (Kielkopf et al. 2001; Selenko et al. 2003; Loerch and Kielkopf 2016). Here we show that in addition to binding U2 snRNA (Yan et al. 1998; Rodgers et al. 2016), Cus2 binds the U2 snRNP through the yeast homolog of SF3b1, Hsh155. A conserved UHM in Cus2 (formerly called RRM2) and a conserved ULM in Hsh155 comprise the major binding interface between the proteins. The Hsh155 ULM interacts with both Cus2 and Tat-SF1 with high affinity *in vitro*. A high-resolution structure of a complex between Tat-SF1 and Hsh155 allows detailed modeling of this interaction from both organisms. Disruption of the Cus2 - Hsh155 interface permits the formation of yeast pre-spliceosomes *in vitro* in the absence of ATP, indicating that ATP is required to remove Cus2 from Hsh155 in order for pre-spliceosomes to assemble. This and other genetic and biochemical data support a model in which Cus2 binds to Hsh155 while refolding U2 snRNA to the stem IIa form necessary for pre-spliceosome assembly, but then must be displaced from Hsh155 by an ATP-dependent activity of Prp5 before stable binding of the U2 snRNP to the pre-mRNA can be established.

## RESULTS

### Identification of a potential UHM-ULM interaction between Cus2 and Hsh155

Primary sequence comparisons identified yeast Cus2 and mammalian Tat-SF1 as UHM-containing proteins (Kielkopf et al. 2001; Kielkopf et al. 2004). Both proteins contain sequences indicating an RRM-like fold, and characteristic residues known to mediate contact with ULMs are found in appropriate positions (Fig. 1A and 1C). In Cus2 these are an acidic residue (D204) important for salt-bridge formation, and a conserved phenylalanine (F235) important for stacking interactions. Although a ULM-containing binding partner of Cus2 has not been identified, Cus2 is reported to interact with the U2 snRNP protein Hsh155 in two-hybrid experiments (Perriman and Ares 2000; Yu et al. 2008; Carrocci et al. 2017). Hsh155 is homologous to human SF3b1 (Pauling et al. 2000), which contains at least five ULMs in its N-terminal sequences (Thickman et al. 2006) (Fig. 1B). Alignment of SF3b1 homologs suggests that Hsh155 contains at least one ULM in its N-terminal sequences (Fig. 1D). Structure guided modeling using previously characterized UHM-ULM complexes (Kielkopf et al. 2004) suggests that Hsh155 and Cus2 share all of the hallmark features of known ULM-UHM interaction partners. The predicted ULM of Hsh155 contains a conserved tryptophan (W101) that could stack with F235 of Cus2 as well as a conserved arginine (R100) that could form the salt bridge with Cus2 D204. Based on this we tested whether these residues are required for an interaction between these proteins.

**Figure 1.**
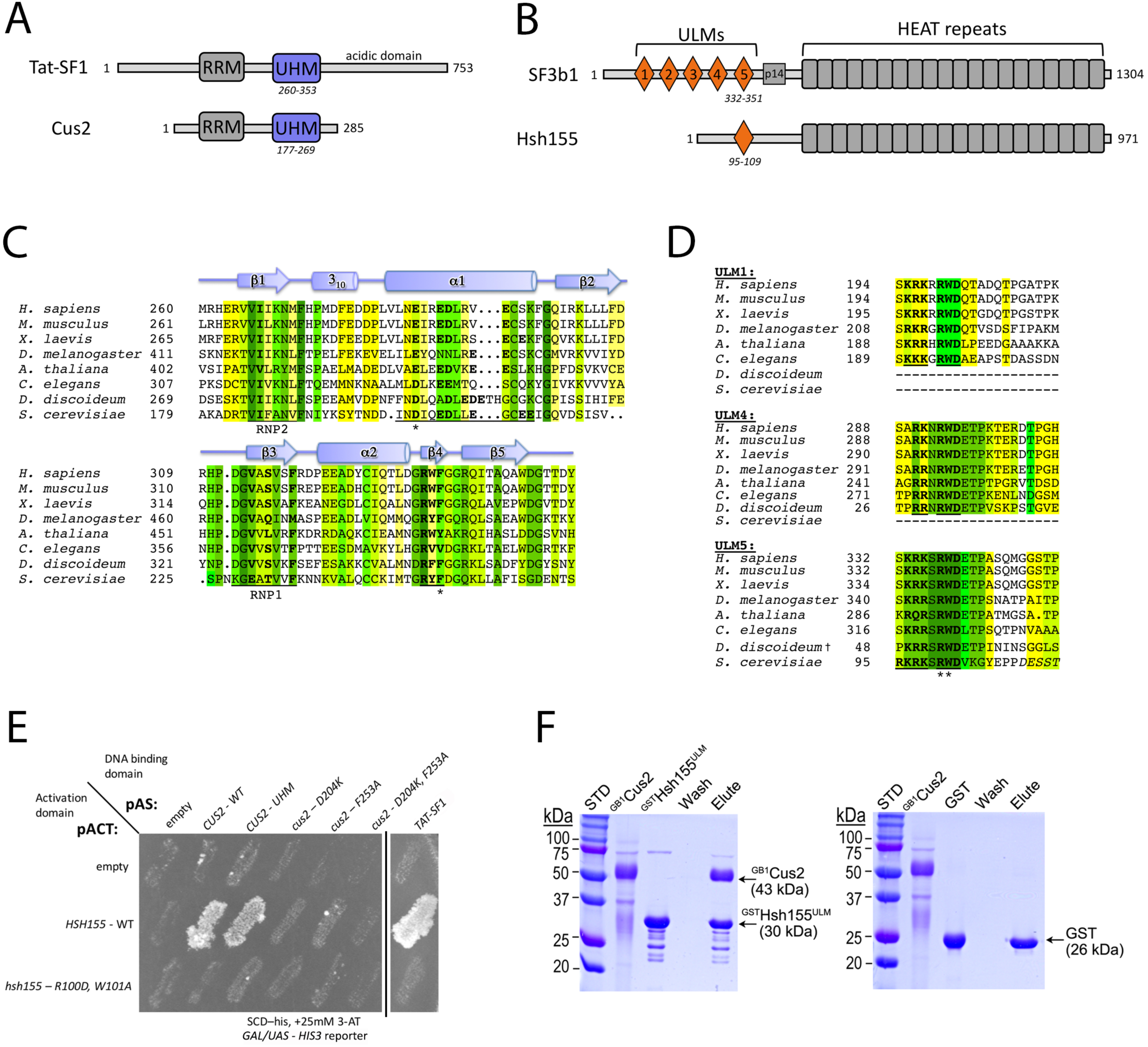
A putative UHM-ULM interaction between the U2 snRNP proteins Cus2 and Hsh155. **(A, B)** Domain organizations of (A) Tat-SF1/Cus2 and (B) SF3b1/Hsh155. **(C**) The sequences of the UHM region of Cus2 (*Saccharomyces cerevisiae*, NCBI RefSeq NP_014113) aligned with homologues including human Tat-SF1 (NCBI RefSeq NP_001156752), *Mus musculus* (NCBI RefSeq NP_083647), *Xenopus laevis* (NCBI RefSeq NP_001083090), *Drosophila melanogaster* (NCBI RefSeq NP_649313), *Arabidopsis thaliana* (NCBI RefSeq NP_197130), *Caenorhabditis elegans* (NCBI RefSeq NP_490765), and *Dictyostelium discoideum* (NCBI RefSeq XP_636172). The residues are colored by identity: <50%, white; 50%, yellow; 63%, chartreuse; 75%, lime; 88% green; 100%, forest. The Cus2 D204 and F253 residues mutated in this study are marked by asterisks. The Tat-SF1 UHM secondary structure elements assigned using the Kabsch and Sander algorithm are indicated schematically above the alignment (Kabsch and Sander 1983). Characteristic sequences of the UHM domain are underlined and bold. The ribonucleoprotein consensus motifs (RNP1 and RNP2) are underlined, and characteristic UHM residues that diverge from the RRM consensus are bold. **(D)** The sequences of the Hsh155 ULM (*S. cerevisiae*, NCBI RefSeq NP_014015) aligned with the ULMs of human SF3b1 (NCBI RefSeq NP_036565) that bind Tat-SF1 (ULM1, ULM4, and ULM5) and homologues including *M. musculus* (NCBI RefSeq NP_112456), *X. laevis* (NCBI RefSeq NP_001084150), *D. melanogaster* (NCBI RefSeq NP_608534), *A. thaliana* (NCBI RefSeq NP_201232), *C. elegans* (NCBI RefSeq NP_497853), and *D. discoideum* (NCBI RefSeq XP_643385). Matching ULMs were identified by Blast (Altschul et al. 1990). The *D. discoideum* homologue appears to contain two rather than five ULMs, and Hsh155 contains a single ULM. Residues are colored by sequence identity as in *C*. Characteristic sequences of the ULM are underlined and bold. For the Hsh155 homologue, residues outside the boundary of the peptide used in this study are italicized, and the bound residues lack regular secondary structure elements. The Hsh155 R100 and W101 residues mutated in this study are marked by asterisks. **(E)** Yeast two-hybrid screen showing interaction between the UHM of Cus2 or Tat-SF1 and the ULM of Hsh155. Sequences encoding wild-type and mutant Cus2 proteins were fused to the DNA binding domain of the *GAL4* transcription factor and tested against constructs expressing either wild-type Hsh155 or mutant Hsh155 proteins fused to the activation domain of *GAL4*. Growth on SCD-His medium supplemented with 25 mM 3-AT indicates interaction. A black vertical line indicates that part of the plate was removed for clarity. **(F)** Cus2 directly and specifically binds the Hsh155 ULM region in GST pull-down assays. The retained proteins were resolved by SDS-PAGE and stained with Coomassie^®^-Brilliant Blue. The sizes of molecular weight standards (STD) and subunits are indicated. GST is a control for nonspecific binding. ^GB1^Cus2 includes an N-terminal GB1-tag to improve solubility and reduce nonspecific binding to the resin. Hsh155^ULM^ includes residues 86-129. The lanes are labeled: ^GB1^Cus2, input Cus2; ^GST^Hsh155ULM or GST, GST or GST-fusion protein used for bait; W, final wash of binding reaction; E, elution.

### Amino acid substitutions in the UHM of Cus2 or the ULM of Hsh155 abolish binding *in vivo*

To test the predicted interaction between Cus2 and Hsh155, we made mutations that substitute key amino acids, and assessed the interaction *in vivo* using the yeast two-hybrid test (Fig. 1E). We observe reporter activation consistent with a protein-protein interaction between full-length, wild-type Hsh155 and full-length Cus2 (Fig. 1E) (Perriman and Ares 2000; Yu et al. 2008; Carrocci et al. 2017). A construct containing the Cus2 UHM alone mediates the interaction. Most importantly, mutations in either the UHM of Cus2 (D204K or F235A) or the ULM of Hsh155 (R100D, W101A) prevent reporter expression, supporting the idea that Cus2 and Hsh155 interact through the predicted UHM and ULM motifs *in vivo*. Tat-SF1, the mammalian homologue of Cus2, also supports a two-hybrid interaction with yeast Hsh155 that is dependent on the ULM of Hsh155, indicating that the interaction is conserved.

To determine whether Hsh155 and Cus2 interact directly, we used glutathione S-transferase (GST) pull-down assays of purified recombinant proteins (Fig. 1F). A protein segment spanning the ULM of Hsh155 (residues 86-129) was fused to GST, immobilized on glutathione-agarose, and incubated with full-length Cus2 protein. The GST-Hsh155 (86-129) protein retained near stoichiometric amounts of Cus2, whereas GST did not detectably bind Cus2, ruling out nonspecific association. Together, these *in vivo* and *in vitro* interaction data indicate that the Cus2 UHM directly and specifically binds to the ULM-region of Hsh155 in a conserved manner.

### The Cus2 UHM binds the Hsh155 ULM with high affinity

We used isothermal titration calorimetry (ITC) to measure the binding affinity of the Hsh155 ULM for full-length Cus2 and homologous Tat-SF1 (residues 1-360 that lacks a divergent C-terminal expansion of acidic residues not found in Cus2) (Fig. 2, Table 1). Cus2 binds the Hsh155 ULM with significant affinity (*K_D_*=32 nM, Fig. 2A and B, Table 1), which is >25-fold greater than the binding affinity of Tat-SF1 for its preferred SF3b1 ULM (ULM5) (Loerch et al. 2018). Tat-SF1 also bound the Hsh155 ULM with subtly higher affinity (*K_D_*=229 nM) than the SF3b1 ULMs (e.g. approximately three-fold compared to ULM5) (Fig. 2A and E, Table 1). This weaker binding of the Tat-SF1 – SF3b1 complex may reflect a more transient, regulated role in humans than the Cus2– Hsh155 complex in yeast, and agrees with differences in the Hsh155 ULM sequence combined with complementary changes in the Cus2 UHM (discussed further below).

**Figure 2.**
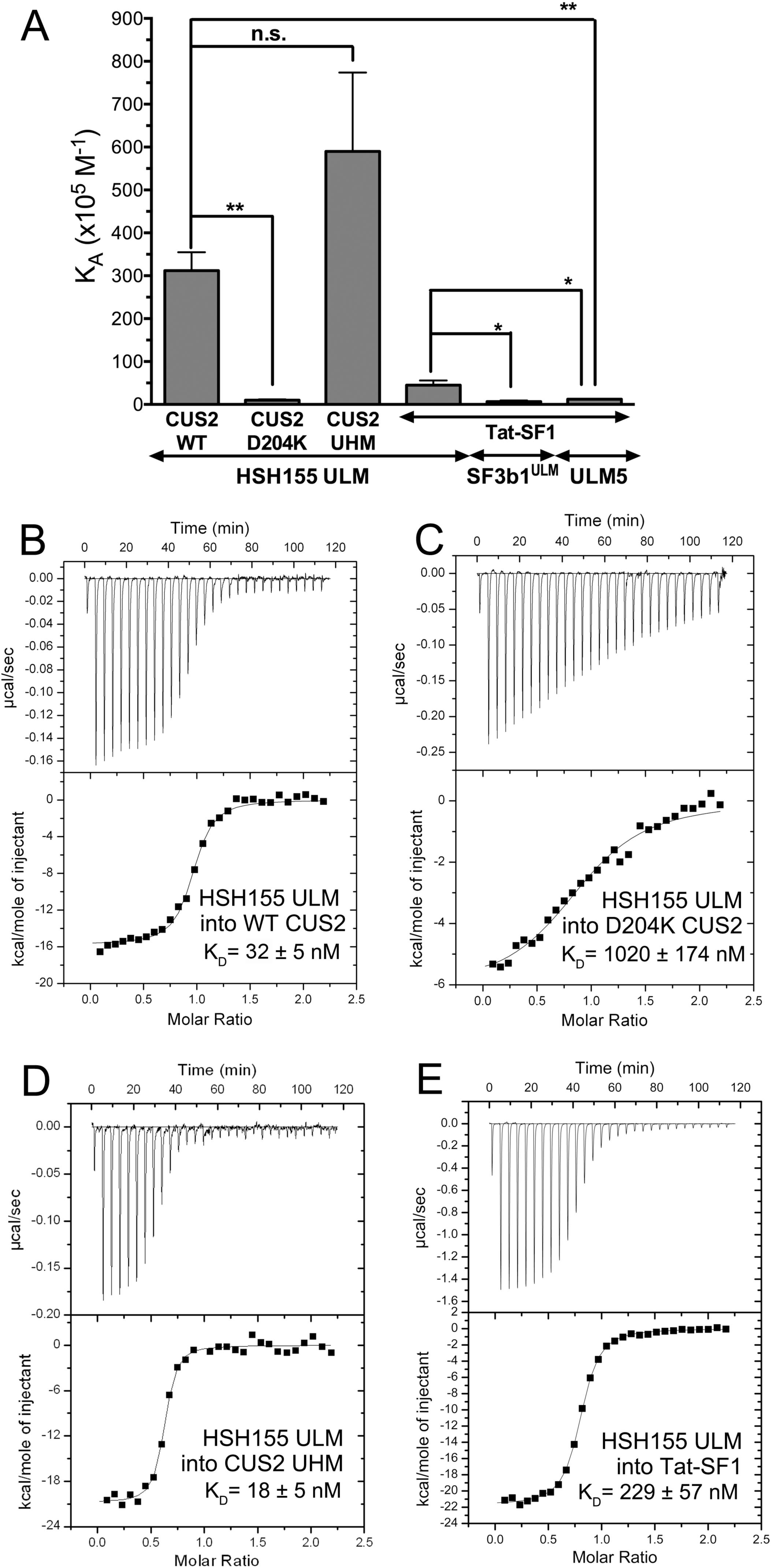
Cus2 and its UHM interact with the ULM of Hsh155 with high affinity in vitro. **(A)** Apparent equilibrium affinities (K_A_) of Hsh155 ULM (residues 95-109) binding to full length Cus2 (isotherm B), D204K-mutant Cus2 (isotherm C), Cus2 UHM (isotherm D, residues 177-269), or Tat-SF1 (isotherm E, residues 1-360). The K_A_’s of the ULM-containing SF3b1 region (SF3b1^ULM^, residues 190-344) or SF3b1 ULM5 (residues 333-351) binding Tat-SF1 were determined under matching conditions (Loerch et al. 2018) and are shown for comparison. Unpaired, two tailed t-tests with Welch’s correction: *, p<0.05; **, p<0.007. (**B** *–* **E**) Representative isotherms of three replicates for each of the indicated pairs of proteins or protein domains. The average fitted K_A_’s and standard deviations are plotted in (A) and given in Table 1.

**Table 1.** Thermodynamics of Tat-SF1 binding SF3b1 ULMs ^a^

| **Interaction:** | **K<sub>D</sub>**<br>(nM) | **ΔG<sup>b</sup>**<br>(kcal mol<sup>-1</sup>) | **ΔH**<br>(kcal mol<sup>-1</sup>) | **-TΔS<sup>b</sup>**<br>(kcal mol<sup>-1</sup>) |
| --- | --- | --- | --- | --- |
| **HSH155 ULM titrated into:** |  |  |  |  |
| WT CUS2 (full length) | 32.4 ± 4.5 | -10.4 ± 0.1 | -16.1 ± 0.5 | 5.7 ± 0.6 |
| D204K CUS2 (full length) | 1020 ± 174 | -6.0 ± 3.0 | -9.4 ± 4.6 | 3.1 ± 1.5 |
| CUS2 UHM | 17.9 ± 4.7 | -10.7 ± 0.2 | -20.9 ± 1.2 | 10.1 ± 1.2 |
| **Tat-SF1 titrated with ULM regions: <sup>c</sup>** |  |  |  |  |
| HSH155 ULM | 229 ± 57 | -9.2 ± 0.1 | -22.7 ± 1.4 | 13.5 ± 1.6 |
| SF3b1<sup>ULM</sup> | 1800 ± 900 | -8.0 ± 0.3 | -66.1 ± 3.0 | 58.0 ± 3.0 |
| ULM5 | 800 ± 100 | -8.4 ± 0.1 | -18.7 ± 1.0 | 10.2 ± 1.2 |

<sup>a</sup> Average values and standard deviations of three independent titrations. <sup>b</sup> Calculated using $\Delta G = -RT \ln (K_D^{-1})$ and $-T\Delta S = \Delta G - \Delta H$ at T=303 K. <sup>c</sup> Reproduced for comparison from Loerch et al. 2018. <sup>d</sup> No heats detected during titration of 200 μM ULM into 20 μM Tat-SF1 protein.

The isolated Cus2 UHM retains the Hsh155 ULM binding affinity (*K_D_*=18 nM) of the full length Cus2 protein (*K_D_*=32 nM), indicating that this domain is responsible for Hsh155 ULM recognition (Fig. 2A and D, Table 1). We further investigated the importance of the Cus2 UHM for binding Hsh155 by mutating a single residue (D204) in the predicted acidic α-helix of the domain. The Cus2 D204K mutation reduces its binding affinity for the Hsh155 ULM by 25-fold (*K*_d_=1020 nM, Fig. 2A and C, Table 1), consistent with the lack of detectable signal for the Cus2 D204K mutant in the yeast two-hybrid screen. Altogether, these data establish that Cus2 forms a high affinity, UHM-dependent complex with the Hsh155 ULM, and that the human homologue Tat-SF1 is sufficiently conserved to support strong binding to the yeast Hsh155 ULM.

### Structural homology of the Cus2 UHM–Hsh155 ULM and Tat-SF1 UHM–Sf3b1 ULM interfaces

To view in detail the interaction between the Hsh155 ULM and one of its binding partner UHMs, we determined a 2.0 Å resolution structure of Tat-SF1 bound to the Hsh155 ULM (residues 98-112) (Figs. 3 and 4, Table 2). Considering its high binding affinity for the Hsh155 ULM and sequence similarity (55%) to the Cus2 UHM, the Tat-SF1 UHM offers a readily-crystallized model system for studying Cus2 UHM-Hsh155 ULM interactions. The Hsh155 ULM is bound to the characteristic α-helical surface of the Tat-SF1 UHM and has well-defined electron density (Fig. 3A). The overall fold and binding site is similar among the structures of the Tat-SF1 UHM bound to Hsh155 or SF3b1 ULMs (r.m.s.d. 0.55 Å between 170 matching UHM atoms or 0.26 Å between 20 matching ULM atoms of this structure and PDB ID 6N3E) (Fig. 3B) (Loerch et al. 2018). The positions of the characteristic UHM and ULM residues are conserved. The Hsh155 R100-W101 residues respectively interact with Tat-SF1 E286 (Cus2 D204) in the acidic UHM α-helix and F337 (Cus2 F253) of the UHM “RXF” motif, in agreement with the importance of these residues for Cus2 – Hsh155 association (Fig. 4A and B). Although the Hsh155 ULM sequence and position in the primary sequence most closely resembles SF3b1 ULM5 (Fig. 1B and D), which has the highest binding affinity for Tat-SF1 among the SF3b1 ULMs, crystal packing prevents the C-terminal residues of SF3b1 ULM5 from interacting with the Tat-SF1 UHM (Loerch et al. 2018). In a different crystal form that avoids such packing contacts, the C-terminal T296-P297 residues of SF3b1 ULM4, which are otherwise identical in sequence to SF3b1 ULM5 (T341-P342), interact with Tat-SF1 W336 at the so-called “X” position of the UHM “RXF” motif (Fig. 4C). In a similar manner as T296-P297 of the SF3b1 ULM4, the C-terminal K104 of the Hsh155 ULM is stacked against the Tat-SF1 W336 side chain (Fig. 4A and C). These consistent conformations of the Hsh155 ULM and SF3b1 ULM4 suggest that an extended interface between the C-terminal ULM residues and the “RXF” motif is a shared feature of the Tat-SF1 and Cus2 UHM.

**Figure 3.**
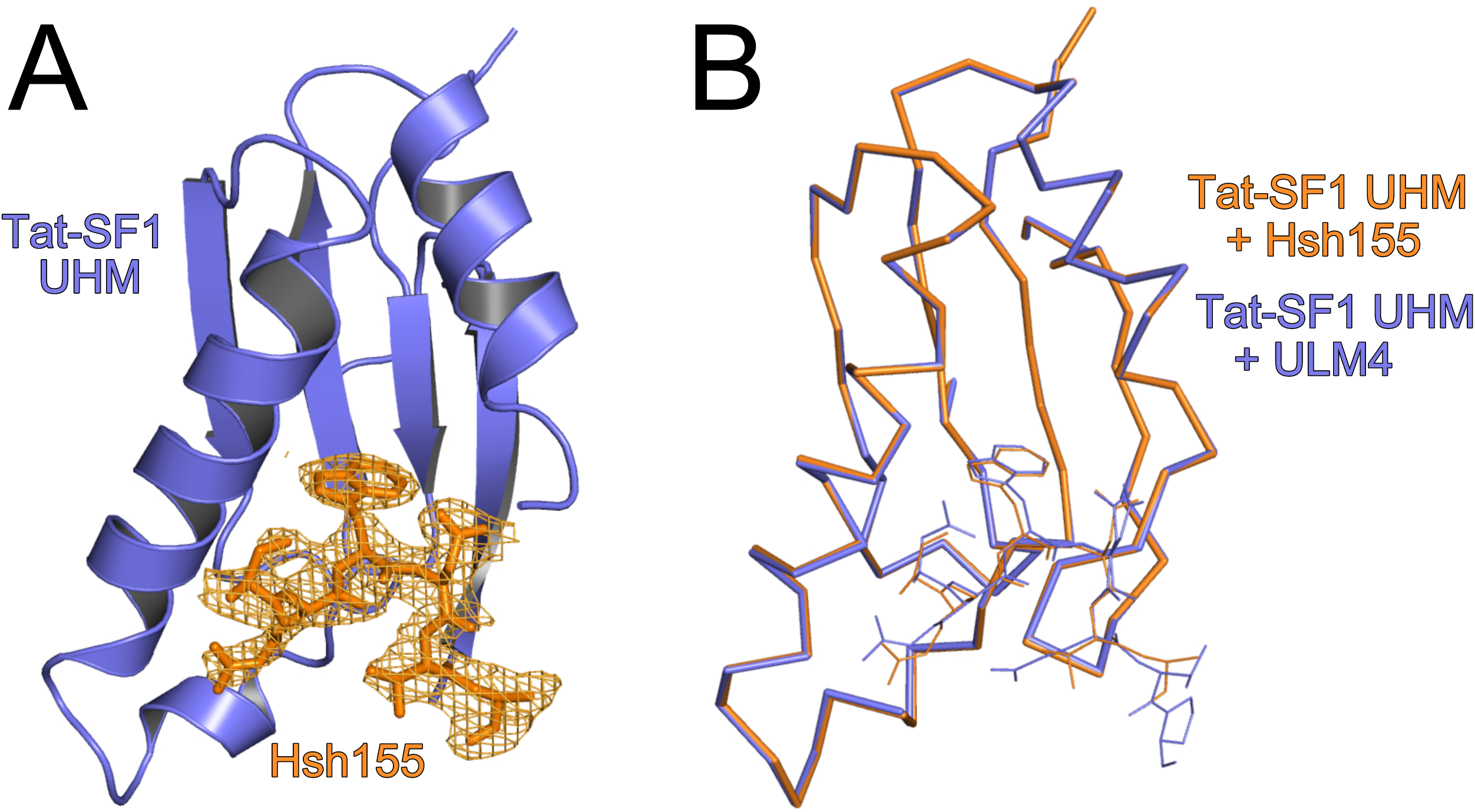
Overall structure of the Hsh155 ULM bound to the Tat-SF1 UHM at 2.0 Å resolution. **(A)** Feature-enhanced electron density map (Afonine et al. 2015) (1.5 *σ* contour level) of Hsh155 ULM (orange) bound to Tat-SF1 UHM (blue). **(B)** Superposed Cα-backbone traces of HSH155-bound compared to ULM4-bound Tat-SF1 UHM structures.

**Figure 4.**
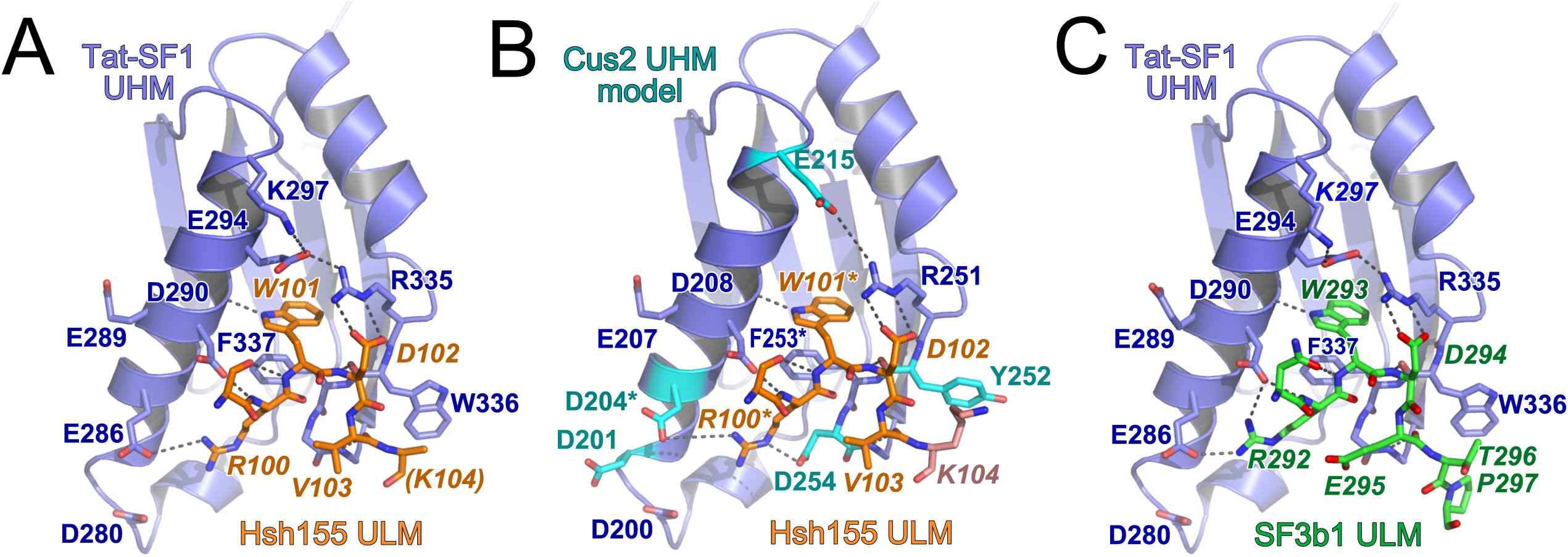
Conserved interfaces revealed by Tat-SF1 – ULM structures. **(A)** Hsh155 ULM (orange) bound to Tat-SF1 UHM (blue). The K104 side chain (parentheses) is disordered and lacks interpretable electron density. **(B)** Model of Hsh155 bound to Cus2 UHM. Interacting residues that differ from the Tat-SF1 homologue were substituted with a sterically compatible rotamer (cyan). Residues modified by mutagenesis (Cus2 D204 and F253; Hsh155 R100 and W101) are marked by asterisks. **(C)** Comparison of human SF3b1 ULM bound to Tat-SF1 UHM (PDB ID 6N3E).

**Table 2.** X-ray data collection and refinement statistics for Tat-SF1 UHM bound to HSH155 ULM. ^a^Values in parentheses refer to the 2.00 – 2.05 Å resolution shells. ^b^For R_free_, 9% of the reflections were randomly-selected then excluded from all stages of refinement.

| **Data collection<sup>a</sup>** |  |
| --- | --- |
| Wavelength (Å) | 0.9795 |
| Resolution range (Å) | 35.16 - 2.00 |
| Space group | P 3<sub>2</sub> 2 1 |
| Unit cell lengths (Å) | 58.58, 58.58, 97.57 |
| Total reflections | 118,606 |
| Unique reflections | 13,598 |
| Multiplicity | 8.7 (8.9) |
| Completeness (%) | 99.9 (99.0) |
| Mean $I/\sigma(I)$ | 12.7 (2.5) |
| $R_{\text{merge}}$ (%) | 8.9 (92.7) |
| $R_{\text{p.i.m.}}$ (%) | 3.2 (32.4) |
| $CC_{1/2}$ | 0.99 (0.88) |
| **Refinement** |  |
| $R_{\text{work}}/R_{\text{free}}$ (%)<sup>b</sup> | 17.0/19.4 |
| No. non-hydrogen atoms | 915 |
| Tat-SF1 UHM | 796 |
| HSH155 ULM | 51 |
| Waters / Chlorides | 66/2 |
| R.m.s. Bonds (Å) | 0.013 |
| R.m.s. Angles (°) | 1.10 |
| Ramachandran (%)<br>favored/allowed/outlier | 100/0/0 |
| Clashscore/percentile (%) | 1.8/99 |
| Avg. B-factor (Å<sup>2</sup>) All | 56.9 |
| Tat-SF1 UHM | 54.3 |
| HSH155 ULM | 99.4 |
| Waters / Chlorides | 55.7/86.7 |
| **PDB Code** | 6NSX |

A Cus2 structure is not yet available for comparison of the Cus2 and Tat-SF1 interfaces with Hsh155. However, *in silico* mutagenesis of the Tat-SF1 UHM suggests structural explanations for the handful of Hsh155 ULM-interacting residues that diverge between these homologues (Fig. 4B). Tat-SF1 E294 is replaced by Cus2 G212, which lacks a side chain to engage the Tat-SF1 R335 (Cus2 R251) in salt bridge covering one face of the ULM tryptophan (Hsh155 W101). Instead, Cus2 E215 in the next turn of the α1-helix is positioned to participate in a similar salt bridge. A Tat-SF1 glycine (G338) following the “RXY” motif is replaced by an aspartate (D254) of Cus2 that could stabilize the position of Hsh155 R100. Notably, glutamates in the SF3b1 ULMs (ULM4 E295 or ULM5 E340) contribute similar negative charges near the corresponding SF3b1 arginine (ULM4 R292 or ULM5 R337), and are replaced by a neutral valine side chain (V103) of the Hsh155 ULM. Near the N-terminus of this helix, a conservative change of a Tat-SF1 glutamate (E286) to aspartate (D204) enables Cus2 to participate in a nearly identical salt-bridge with Hsh155 R100. A nearby Cus2 aspartate (D201) is likely to account for the additional arginine (R95) at the N-terminus of the Hsh155 ULM, which is disordered in the Tat-SF1-bound structure. Lastly, the substitution of Cus2 Y252 for Tat-SF1 W336 in the “RXF” motif is accompanied by a reciprocal change of the interacting SF3b1 ULM threonines (ULM4 T296 or ULM5 T341) to Hsh155 K104. This Hsh155 lysine is capable of donating a hydrogen bond to the hydroxyl group of the Cus2 tyrosine, whereas the Tat-SF1 tryptophan offers a hydrogen bond donor from the imine-nitrogen. Accordingly, the Hsh155 K104 side chain is disordered and lacks clear electron density in the Tat-SF1 UHM complex. The corresponding SF3b1 threonines are highly phosphorylated in humans (Wang et al. 1998; Bessonov et al. 2010; Agafonov et al. 2011; Girard et al. 2012), such that the divergent replacement with lysine in yeast (Fig. 1D) may also reflect distinct regulation.

Altogether, we conclude that the ULM of Hsh155 binds to the Tat-SF1 UHM in the characteristic fashion observed for other examples of this protein interaction motif, and the structural similarity between Tat-SF1 and Cus2 provides a detailed model for the molecular basis of Cus2 UHM binding to the ULM of Hsh155.

### Disruption of the Hsh155-Cus2 interaction does not perturb growth or splicing catalysis *in vitro*

To understand the importance of the high-affinity binding between Cus2 and Hsh155 for spliceosome assembly and catalysis, we created a mutant yeast strain in which key residues of the Hsh155 ULM (R100D, W101A or “RW-DA”, Fig. 1D) are altered to block Cus2 binding. Since deletion of *CUS2* is not lethal (Yan et al. 1998), we anticipated that the loss of its binding to Hsh155 would also not be lethal, and a recent analysis of Hsh155 deletions in this region is consistent with this (Carrocci et al. 2018). Successful CRISPR/Cas9 editing of the RW-DA mutation into the chromosomal copy of *HSH155* creates a viable strain with no obvious growth defect at a variety of different temperatures (Supplemental Fig. S1B). Furthermore, a tagged version of the Hsh155^RW-DA^ protein is expressed at wild-type levels as judged by western blotting (Supplemental Fig. S1A).

To evaluate pre-mRNA splicing in the Hsh155^RW-DA^ mutant, we generated cell free extracts and performed *in vitro* splicing reactions. In our experience, yeast splicing mutants with undetectable *in vivo* growth defects can show clear and informative spliceosome assembly and splicing defects *in vitro*. Nonetheless, splicing catalysis of exogenous actin pre-mRNA substrate in the Hsh155^RW-DA^ mutant extract was indistinguishable from wild type in ATP dependence (Fig. 5A, lanes 1 and 3), and extent (Fig. 5A, lanes 2 and 4). This indicates that disrupting the interaction between Hsh155 and Cus2 does not greatly impact the overall chemistry of splicing.

**Figure 5:**
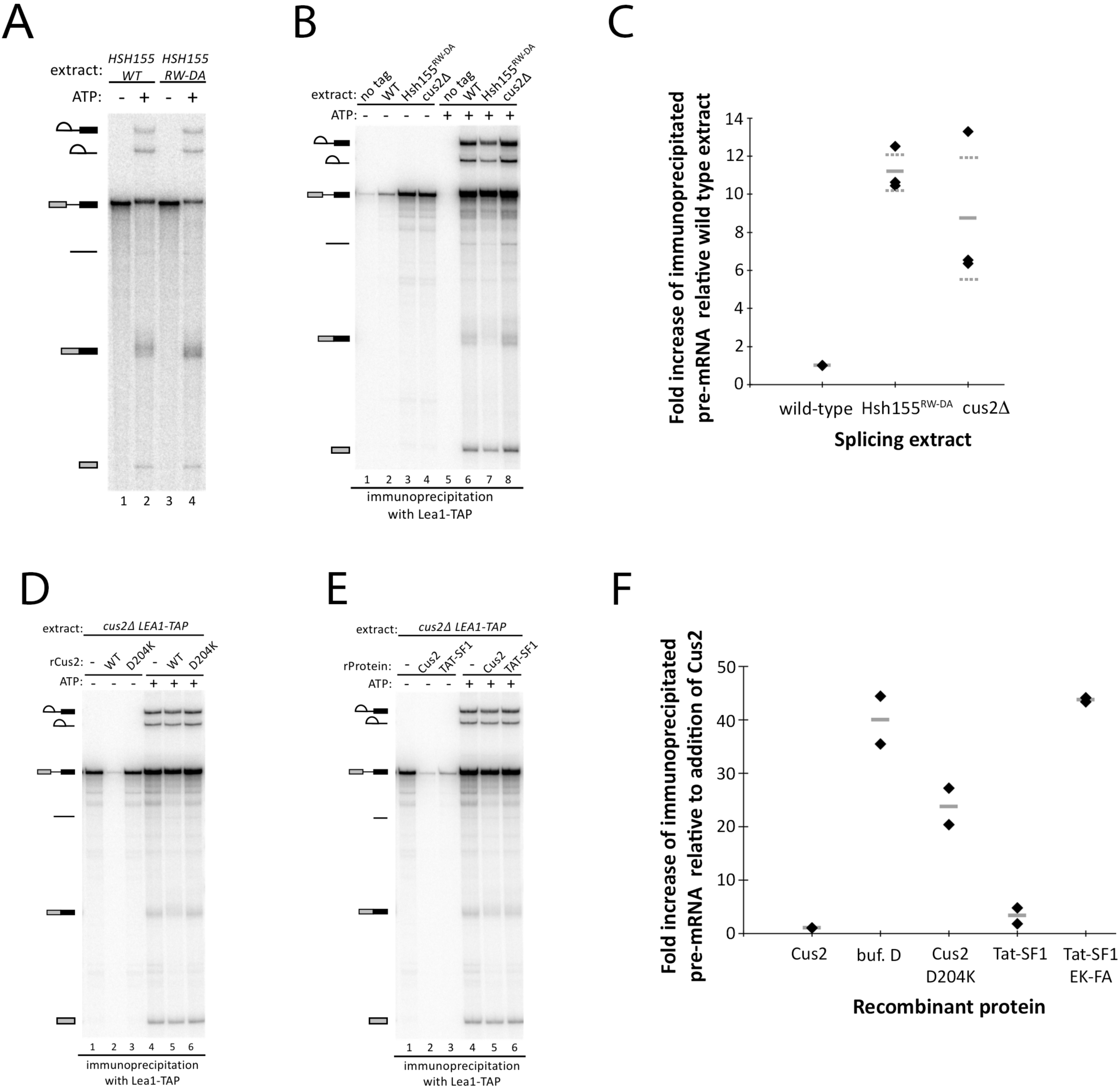
The interaction between the UHM of Cus2 and the ULM of Hsh155 is functionally conserved, and disruption of the interaction permits pre-spliceosome assembly in the absence of ATP. **(A)** In vitro analysis of splicing of radiolabeled actin pre-mRNA. Pre-mRNA was incubated with either wild-type or Hsh155^RW-DA^ splicing extracts for 20 minutes with or without ATP. RNA was extracted from the reactions, resolved on a 6% acrylamide/8M urea gel, and visualized by autoradiography. Precursor, intermediates, and products are indicated. **(B)** Co-immunoprecipitation of splicing complexes with Lea1-TAP in wild-type, Hsh155^RW-DA^, and cus2Δ extracts in the presence or absence of ATP. Reactions were set up as described in (A). Following the reaction, Lea1-TAP-containing splicing complexes were immunoprecipitated with IgG-conjugated magnetic beads. After extensive washing of the beads, RNA was extracted, resolved on a denaturing acrylamide gel, and visualized with autoradiography. Co-immunoprecipitation of pre-mRNA in the absence of ATP serves as a proxy for ATP-independent pre-spliceosome formation. A “no tag” extract serves as a control for background binding of splicing complexes to the IgG-conjugated beads. The autoradiograph shown is representative of triplicate experiments. **(C)** Quantification of triplicate experiments described in (B) indicating the fold increase of immunoprecipitated pre-mRNA by Lea1-TAP from cus2Δ and Hsh155^RW-DA^ extracts relative to the wild-type extract in the absence of ATP. Data points are indicated by diamonds, the mean is indicated by a solid gray line, and the standard deviation of the mean is indicated by dashed gray lines. **(D)** Reenforcement of ATP-dependent pre-spliceosome assembly by addition of recombinant Cus2 proteins. Cus2Δ Lea1-TAP extract was depleted of ATP in the presence of either buffer D or 500 nM wild-type or D204K mutant recombinant Cus2. Pre-mRNA and either water or ATP was then added to the extract and splicing was allowed to proceed for 20 minutes. Splicing complexes were purified and co-immunoprecipitated RNA was extracted and visualized as described in (B). **(E)** Reenforcement of ATP-dependent pre-spliceosome formation by addition of recombinant Cus2 or Tat-SF1. Cus2Δ Lea1-TAP extract was depleted of endogenous ATP in the presence of either buffer D or 500 nM recombinant Cus2 or 1 µM Tat-SF1. Pre-mRNA and either water or ATP was then added to the reactions and splicing was performed for 20 minutes. Splicing complexes were purified and co-immunoprecipitated RNA was extracted and visualized as described in Figure (B). Autoradiographs shown in (D) and (E) are representative of duplicate experiments. **(F)** Quantification of duplicate experiments show in (D) and (E) indicating the fold increase of ATP-independent immunoprecipitated pre-mRNA by Lea1-TAP from reconstituted splicing reactions relative to the addition of recombinant wild-type Cus2. Data points are indicated by diamonds and the mean is indicated by a solid gray line.

### Disruption of the Hsh155-Cus2 interaction bypasses the ATP requirement for pre-spliceosome formation

Stable association of the U2 snRNP with the pre-mRNA BP in the pre-spliceosome is dependent on ATP and Prp5 (Liao et al. 1992; Ruby et al. 1993; Perriman et al. 2003). Previous studies have shown that this ATP-dependence can be bypassed under several different genetic and biochemical conditions (Table 3), including disrupting or deleting the Cus2 protein (Perriman and Ares 2000; Perriman et al. 2003; Macias et al. 2008; Liang and Cheng 2015), hyperstabilizing the stem IIa conformation of U2 (Perriman et al. 2003; Perriman and Ares 2007), or destabilizing the BP interacting stem loop (BSL) of U2 that must open to form the extended duplex with the intron observed in the pre-spliceosome and the later complexes (Perriman and Ares 2010). A consistent interpretation is that Prp5 normally binds ATP to displace Cus2 from the U2 snRNP so that stable binding of U2 snRNP to the pre-mRNA can occur.

**Table 3:**
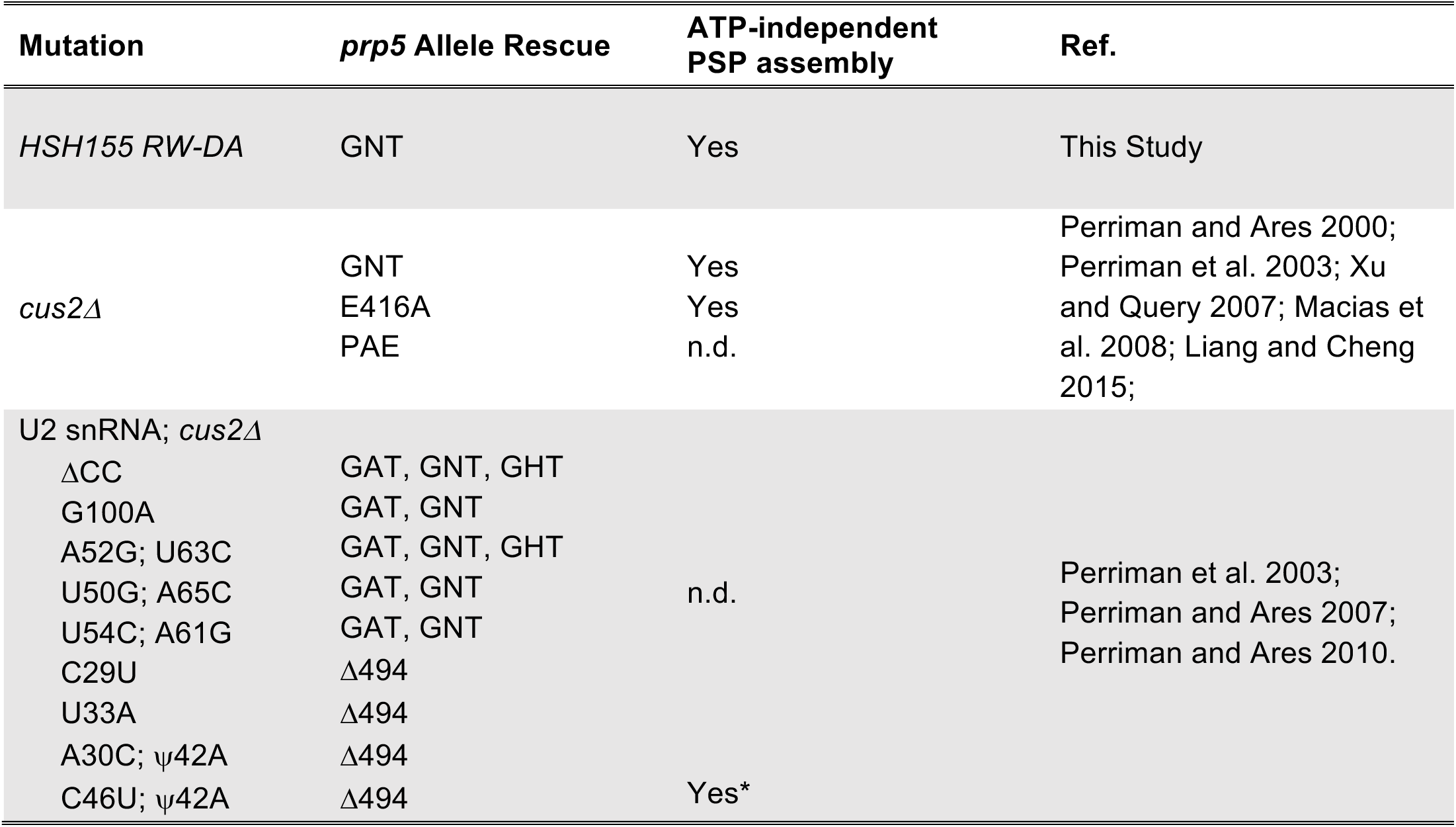
Mutations that bypass the ATP-dependent function of Prp5 during stable base pairing of the U2 snRNP with the pre-mRNA BP.

To test whether the binding of the Cus2 UHM to the ULM of Hsh155 is required to enforce the ATP-dependence for stable U2 snRNP binding to pre-mRNA, we generated yeast strains in which the U2 snRNP protein Lea1 was fused to the TAP-tag at its C-terminus and introduced the RW-DA ULM mutation into *HSH155* in the *LEA1-TAP* strain by CRISPR/Cas9 editing (DiCarlo et al. 2013). As a control for ATP-independent pre-spliceosome assembly, we created a second *LEA1-TAP* strain deleted for the *CUS2* gene. We then tested splicing extracts from these strains for the ability to assemble stable U2 snRNP-pre-mRNA complexes under different conditions by immunoprecipitation using the TAP-tag on Lea1 (Fig. 5B-F).

Under splicing conditions in the presence of ATP, the U2 snRNP associates with pre-mRNA and all splicing intermediates and products, except released spliced exons. And indeed Lea1-TAP co-immunoprecipitates radiolabeled pre-mRNA, splicing intermediates, and products, including spliced exons likely still associated with the spliceosome. Similar levels of U2-bound pre-mRNA are observed in the wild-type, *hsh155^RW-DA^*, and *cus2Δ* extracts, as compared to an untagged extract (Fig. 5B, lanes 5-8; Supplemental Fig. S2A, lanes 5-8). When ATP is absent from the reactions, we observe little to no radiolabeled pre-mRNA co-immunoprecipitating in wild-type extracts, consistent with the ATP requirement for stable binding of U2 to pre-mRNA in pre-spliceosomes (Fig. 5B, lanes 1 and 2 and 5C; Supplemental Fig. S2A, lanes 1 and 2). Remarkably, extracts expressing the Hsh155^RW-DA^ protein produce significant amounts of U2-associated pre-mRNA complexes in the absence of ATP (∼11 fold more compared to a wild-type extract), similar to that observed when Cus2 is absent from the extract (∼8 fold more than a wild-type extract, Fig 5B, lanes 3 and 4 and 5C; Supplemental Fig. S2A, lanes 3 and 4 and S2G, lane 2). In an orthogonal approach, we assayed the formation of splicing complexes by native agarose gel electrophoresis either in the presence or absence of ATP, or the presence of non-hydrolyzable AMP-PCP. For all extracts tested, addition of ATP to the reactions results in the formation of pre-spliceosomes and spliceosomes within 20 minutes (Supplemental Fig. S2B, lanes 1, 4, and 7). Most importantly, whereas wild-type extracts fail to form pre-spliceosomes in the absence of ATP or in the presence of AMP-PCP, the Hsh155^RW-DA^ extracts do so at levels comparable to those observed when Cus2 is disrupted (Supplemental Fig. S2B, compare lanes 2 and 3 with 5, 6, 8, and 9). We conclude that binding of Cus2 to the ULM of Hsh155 enforces the ATP requirement for pre-spliceosome formation *in vitro*, and this requirement can be bypassed either when Cus2 is absent in the extract, or when Cus2 is present but unable to bind to Hsh155 (Fig. 5B and C).

### Reconstitution of ATP-dependence requires a functional Cus2 UHM-Hsh155 ULM interaction

The ATP-dependence of pre-spliceosome formation that is lost in extracts lacking Cus2 can be restored by addition of recombinant Cus2 (Perriman and Ares 2000). To test whether enforcement of ATP-dependence requires the ability of Cus2 to bind Hsh155 we reconstituted *cus2Δ LEA1-TAP* splicing extracts with either recombinant wild-type Cus2 or the Cus2^D204K^ UHM mutant, which binds only poorly to Hsh155 (Fig. 1E; Fig. 2A and C; Table 1). After reconstitution and splicing were complete, U2 snRNP-pre-mRNA complexes were immunoprecipitated using the TAP-tag on Lea1, and the amount of bound RNA was determined on denaturing acrylamide gels (Fig. 5D and F; Supplemental Fig. S2C and G). In the absence of ATP and without addition of Cus2, pre-spliceosomes form in the extract, judged as before by pre-mRNA binding with the U2 snRNP (Figs. 5D, lane 1 and 5F; Supplemental Fig. S2C, lane 1 and S2G, lane 2). As shown previously (Perriman and Ares 2000), addition of recombinant wild-type Cus2 restores the requirement of ATP for pre-spliceosome formation, as little to no pre-mRNA is co-immunoprecipitated with Lea1-TAP (Figs. 5D, lane 2 and 5F; Supplemental Fig. S2C, lane 2 and S2G, lane 3). Interestingly, addition of recombinant Cus2^D204K^ partially restores the requirement of ATP suggesting this protein may retain a small amount of Hsh155 ULM binding activity under *in vitro* splicing conditions. However, when Cus2^D204K^ is added to the reactions at the same concentration as wild-type Cus2 it is not sufficient to fully enforce the ATP requirement, as shown by formation of pre-spliceosomes at levels intermediate to those observed with or without added wild-type Cus2 (Fig. 5D, lane 3 and 5F; Supplemental Fig. S2C, lane 3 and S2G, lane 4). The addition of recombinant Cus2 proteins does not interfere with splicing in the presence of ATP (Fig. 5D, compare lane 4 to 5 and 6; Supplemental Fig. S2C, lanes 4-6), indicating the inhibition of pre-spliceosome formation in the absence of ATP is not due to a general negative effect of adding recombinant protein to the extract. These results parallel those observed when native gel electrophoresis is used to monitor splicing complex formation (Supplemental Fig. S2D). We conclude that Cus2 must be able to bind to the ULM of Hsh155 with high affinity in order to enforce the ATP-dependence of pre-spliceosome formation.

Human Tat-SF1 binds the yeast Hsh155 ULM with affinity that approaches that of Cus2 (within seven-fold; Fig. 2A and E; Table 1), and we wondered whether it could enforce the ATP requirement for pre-spliceosome formation in yeast extracts. Like the addition of Cus2, addition of recombinant Tat-SF1 (residues 1-360, lacking the divergent C-terminus) restores the ATP-requirement for stable binding of the U2 snRNP with pre-mRNA as judged by the greatly reduced levels of U2 snRNP-bound pre-mRNA when Tat-SF1 is added compared to the buffer only control (Fig. 5E, lanes 1-3 and 5F; Supplemental Fig. S2E, lanes 1-3 and S2G, lanes 2,3, 5-10). However, reenforcement of ATP dependent pre-spliceosome formation required two-fold more Tat-SF1 (1 µM) in the splicing reactions compared to Cus2 (500 nM). Furthermore, addition of recombinant Tat-SF1 with mutations in the UHM at amino acids important for SF3b1 ULM binding (E286K/F337A, Loerch et al. 2018) does not restore the requirement of ATP for pre-spliceosome formation (Fig. 5F; Supplemental Fig. S2G, lanes 11-13). Addition of Tat-SF1 to splicing reactions does not inhibit splicing or the formation of U2 snRNP-containing splicing complexes in the presence of ATP (Fig. 5E, lane 6 and Supplemental Fig. S2E, lane 6). Similar results are observed when complexes are evaluated by native gel electrophoresis (Supplemental Fig. S2F). We conclude that human Tat-SF1 is sufficiently structurally related to Cus2, in particular with respect to its interaction with the Hsh155 ULM, that it can functionally replace Cus2 for enforcement of the ATP requirement for pre-spliceosome formation *in vitro*.

### Mutation of the Hsh155 ULM suppresses a lethal Prp5 ATP binding site mutation

Previous studies have shown that an ATP-dependent function of Prp5 is dispensable when Cus2 is absent, either for growth *in vivo* or for splicing *in vitro* (Table 3) (Perriman et al. 2003; Liang and Cheng 2015). Since Cus2-containing extracts from cells mutated for the Cus2 binding site on Hsh155 form pre-spliceosomes without ATP *in vitro* (Fig. 5), we wondered whether the mutation in the ULM of Hsh155 would suppress the lethality of a *prp5* ATP-binding mutant in cells expressing wild-type Cus2. We introduced the Hsh155^RW-DA^ mutations by editing a yeast strain permissive for testing *prp5* mutants by plasmid shuffling (chromosomal deletion of *PRP5* with wild-type *PRP5* on a *URA3* plasmid) (Perriman et al. 2003). Two such strains expressing either wild-type or ULM-mutant *hsh155* were transformed with a second plasmid carrying either wild-type *PRP5* or mutant *prp5-GNT*, which has a K to N substitution in the highly conserved GKT motif responsible for binding ATP. No viable colonies are observed after selection for cells that spontaneously lose the plasmid borne wild type *PRP5* gene when cells have wild type *HSH155* and *prp5-GNT* as the only copy of *PRP5*, confirming that the *prp5*-*GNT* mutation is lethal in a wild type background (Fig. 6, right). In contrast cells expressing HSH155^RW-DA^ are viable with *prp5-GNT*, indicating that disrupting the binding of Cus2 to Hsh155 suppresses this Prp5 defect. We conclude that in the absence of Cus2 binding to Hsh155, the ATP-binding function(s) of Prp5 are dispensable *in vivo* (Fig. 6, right).

**Figure 6:**
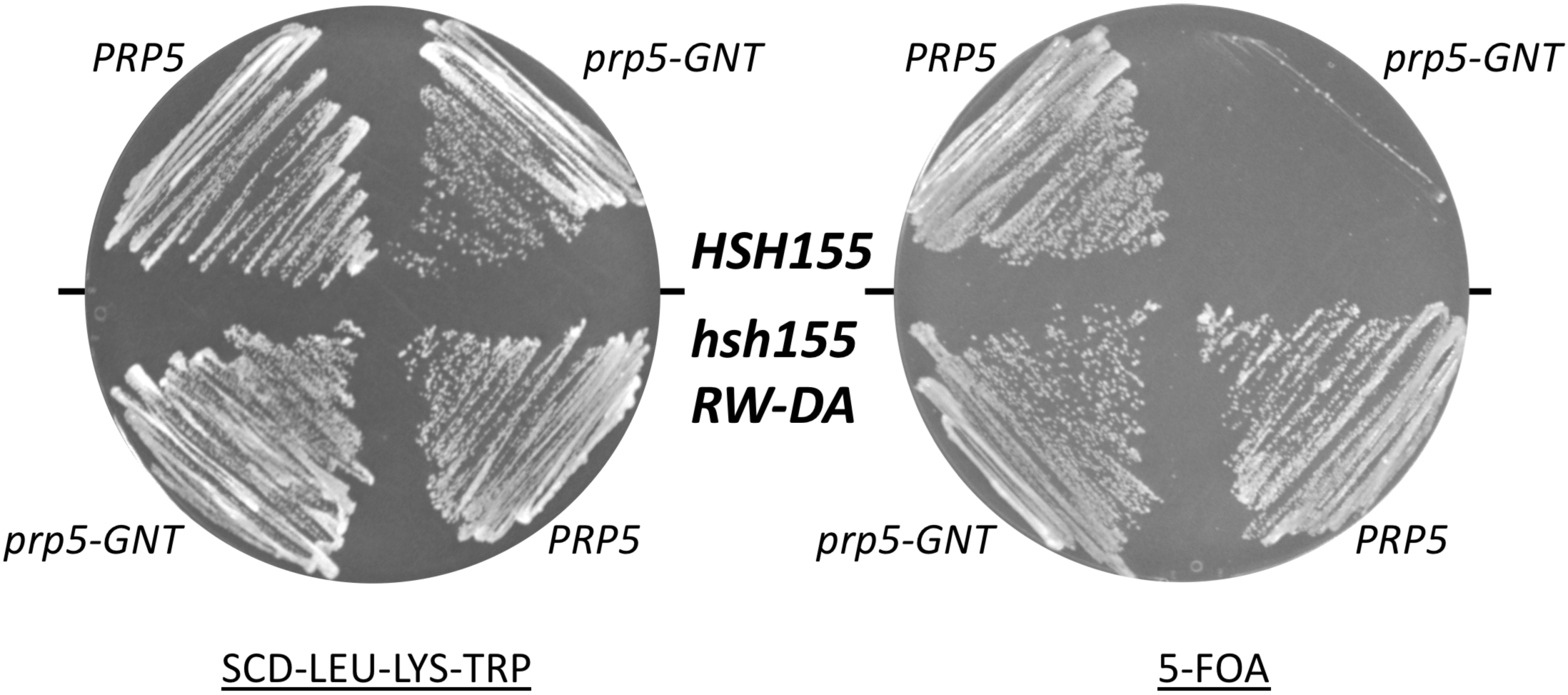
Disruption of the interaction between the UHM of Cus2 and the ULM of Hsh155 bypasses the essential ATP-dependent function of Prp5. The ULM of Hsh155 was mutated (R100D, W101A) in yeast strain DS4D (*prp5Δ, cus2Δ, u2Δ*; Perriman et al., 2003) harboring pRS317-*CUS2*, pRS315-*U2*, and pRS316-*PRP5*. The wild-type and *hsh155^RW-DA^* mutant strains were then transformed with pRS314-*prp5-GNT* (left) and then plated on medium containing 5-FOA (right) to select for cells able to lose the pRS316-*PRP5* plasmid.

## DISCUSSION

Here we show that Cus2 and its human homolog Tat-SF1 bind the core U2 snRNP protein Hsh155 (human homolog SF3b1) through conserved UHM and ULM domain surfaces (Figs. 1-4) *in vivo* (Fig. 1E) and *in vitro* with high affinity (Figs. 1F, 2). Mutations in the UHM of Cus2 or the ULM of Hsh155 disrupt the binding (Figs. 1E and 2, Table 1), altering the function of these proteins. A crystal structure of the complex between Tat-SF1 and Hsh155 solved at 2.0 Å resolution (Fig. 3) reveals the molecular details and demonstrates the evolutionary conservation of this binding interaction (Fig. 4). Importantly, biochemical and genetic assays show that binding between Cus2 and Hsh155 enforces the ATP-dependence of pre-spliceosome assembly (Figs. 5, S2). Mutations that disrupt binding between Cus2 and Hsh155 permit pre-spliceosome assembly in the absence of ATP *in vitro* (Fig. 5) and suppress the lethality of a Prp5 ATP-binding mutant *in vivo* (Fig. 6). Together these data support a model in which Prp5 ATP binding or hydrolysis releases Cus2 from the ULM of Hsh155, allowing the stable base pairing between U2 snRNA and the intron BP that is a defining characteristic of the A-complex (Fig. 7).

**Figure 7:**
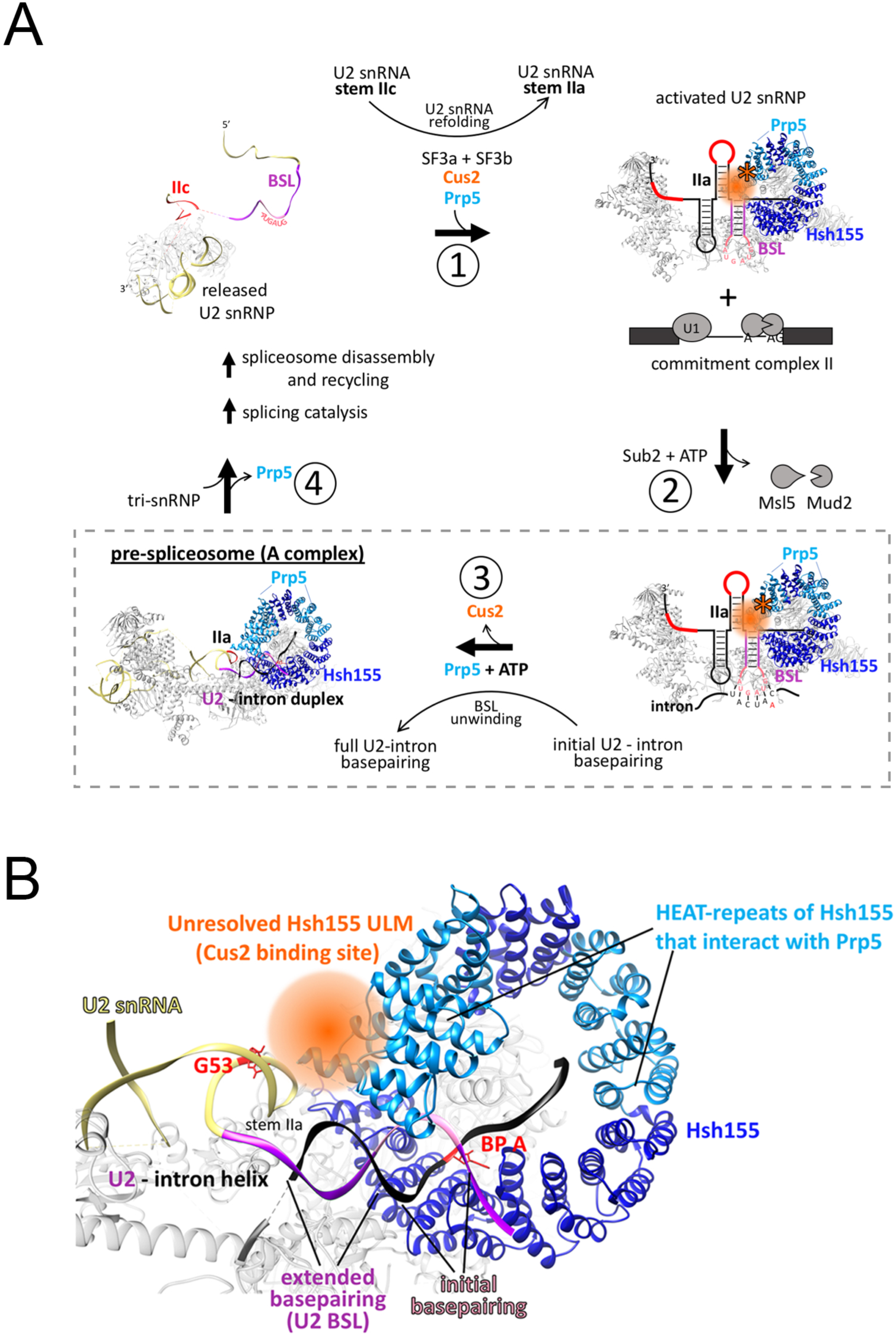
A model of ATP-dependent prespliceosome assembly. **(A)** After a completed round of splicing and spliceosome disassembly, U2 snRNA is released from the spliceosome in the stem IIc (red) form and lacks the SF3a and SF3b subcomplexes. (1) Prior to another round of splicing, the BSL (purple) of U2 snRNA is reformed and Cus2 (orange) associates with both U2 snRNA and the ULM of Hsh155 to promote the transition from stem IIa to stem IIc. (2) The activated U2 snRNP then engages the intron BP in the commitment complex through a limited number of initial base pairs between the BP and the loop of the BSL (pink), possibly displacing the Msl5-Mud2 heterodimer from the commitment complex. (3) ATP hydrolysis by Prp5 disrupts the Cus2-UHM/Hsh155 ULM interaction to release Cus2 and unwinds the helix of the BSL to promote stable duplex formation between U2 snRNA and the intron BP generating the prespliceosome. (4) Stable duplex formation between U2 snRNA and the intron BP results in release of Prp5 and recruitment of the U4/U5/U6 tri-snRNP. Aspects of this model specific to this work are boxed with a dashed line. **(B)** Cryo-EM model (PDB 6G90) of the yeast pre-spliceosome showing the putative binding site of Cus2 relative to Hsh155, the U2/intron helix, and stem IIa of U2 snRNA. The unresolved ULM of Hsh155 and the space likely occupied by Cus2 is orange. HEAT-repeats of Hsh155 previously shown to interact with Prp5 are light blue. Sequences of U2 snRNA that compose the BSL are purple (stem) and pink (loop). The G53A mutation in U2 snRNA that destabilizes stem IIa and is suppressed by Cus2 is indicated in red.

### Key steps in pre-spliceosome assembly and the role of ATP

During the RNA rearrangements associated with the first catalytic step of splicing, U2 snRNA is remodeled to disrupt stem IIa and form stem IIc (Hilliker et al. 2007; Perriman and Ares 2007; Galej et al. 2016; Rauhut et al. 2016; Wan et al. 2016; Yan et al. 2016; Bai et al. 2017; Fica et al. 2017; Liu et al. 2017; Plaschka et al. 2017; Wan et al. 2017; Wilkinson et al. 2017; Yan et al. 2017; Plaschka et al. 2018). However, before it can be reused in a subsequent round of splicing U2 snRNA must be refolded into the stem IIa form, which makes critical contacts with SF3a and SF3b proteins for pre-spliceosome assembly (Liu et al. 2017; Wilkinson et al. 2017; Plaschka et al. 2018). Based on its genetic interactions with stem IIa mutants (Yan et al. 1998) and its effect on U2 snRNA structure in single molecule studies (Rodgers et al. 2016), Cus2 plays an important but not essential role in refolding U2 snRNA. The BSL, a highly conserved U2 snRNA stem-loop immediately upstream of stem IIa, is inferred from genetic studies to present bases 34-38 to nucleate pairing with the intron BP (Perriman and Ares 2010). Unwinding the BSL is necessary to form the extended U2-intron duplex in the pre-spliceosome, and may depend on Prp5, based on genetic interactions (Perriman and Ares 2010) and UV-crosslinking results (Liang and Cheng 2015).

ATP hydrolysis is required for pre-spliceosome assembly, and mutations of Prp5 that alter its ATP binding and hydrolysis activity cannot support pre-spliceosome assembly *in vitro* (Perriman et al. 2003). Previous results show that in the absence of Cus2, lethal Prp5 mutants incapable of ATP binding are viable, indicating that loss of Cus2 bypasses the need for ATP *in vivo* and *in vitro* (Perriman et al. 2003; Xu and Query 2007). Here we show that this same function of Prp5 is unnecessary when Cus2 is present but cannot bind Hsh155 (Fig. 6). Given the location of the binding site for Cus2 between the unresolved N-terminus of Hsh155 and the extended U2-intron duplex in A-complex (Fig. 7B) (Plaschka et al. 2018), it seems clear that ATP-dependent displacement of Cus2 by Prp5 is a critical role of ATP in pre-spliceosome formation (Fig. 7A). Thus, although it is not itself essential, removal of Cus2 from Hsh155 represents the essential ATP-dependent function of Prp5.

Other RNA helicase family members including Prp5 have been implicated in fidelity checkpoints at different steps in the splicing pathway (Burgess and Guthrie 1993; Mayas et al. 2006; Xu and Query 2007; Koodathingal et al. 2010; Tseng et al. 2011; Yang et al. 2013; Wlodaver and Staley 2014; Liang and Cheng 2015). Outstanding questions remain concerning how the U2 snRNP discriminates between correct and incorrect intron BPs, and whether Cus2 plays a role in fidelity given its relationship to the ATP-hydrolysis activity of Prp5. Query and colleagues have proposed a Prp5-mediated fidelity event in which a conformational change due to Prp5 ATP hydrolysis promotes a discard pathway for incorrect intron BPs that is in kinetic competition with U2 snRNA base pairing with a correct intron BP (Xu and Query 2007). However, more recent work from the Cheng group suggests that Prp5 exerts a fidelity check that is independent of ATPase activity, whereby formation of the correct U2-intron duplex lowers the affinity of Prp5 for the pre-spliceosome, allowing recruitment of the tri-snRNP (Liang and Cheng 2015). Formation of an incorrect U2-intron duplex results in retention of Prp5, which presents a block to tri-snRNP recruitment, and presumably triggers an as yet uncharacterized disassembly pathway. Mutations of Prp5 that enhance or decrease ATP-hydrolysis appear to promote the onward use of incorrect BPs in this assay by more readily dissociating from incorrectly assembled pre-spliceosomes, revealing a lack of correspondence between the rate of ATP-hydrolysis and the effectiveness of the fidelity check (Liang and Cheng 2015). Since the fidelity check revealed by this assay appears indifferent to both ATP-hydrolysis and the presence of Cus2, it seems unlikely that Cus2 makes a direct contribution to the fidelity of BP selection, except through its general antagonism of Prp5 function (Xu and Query 2007).

Based on these observations, we propose a model whereby Cus2, through binding to the stem IIc form of U2 snRNA and the ULM of Hsh155, assists in refolding U2 snRNA into the stem IIa form as the SF3b and SF3a proteins join the U2 snRNP (Fig. 7A, i). The BSL of U2 snRNA engages the pre-mRNA BP within the commitment (or E-) complex through a limited number of initial base pairs, possibly displacing or destabilizing the Msl5-Mud2 heterodimer which is bound at the intron BP in the commitment complex (Fig. 7A, ii). In order to generate the extended duplex between U2 snRNA and the intron BP, the ATP-hydrolysis function of Prp5 must remove Cus2 from the ULM of Hsh155, so that the BSL can unwind to establish the U2 – intron duplex, which is bound by Hsh155 to stabilize the pre-spliceosome (Fig. 7A, iii). The exact timing and relationships between Prp5 ATP-hydrolysis, release of Cus2 from Hsh155, unwinding of the BSL and insertion into Hsh155 remain unknown. Formation of a correct U2 – intron duplex successfully bound into Hsh155 results in the release of Prp5 and progression of the splicing pathway (Fig. 7A, iv).

### Insights into human pre-spliceosome assembly

Given the greater diversity of pre-mRNA BP sequences used by mammals compared to yeast, it is still unclear whether Tat-SF1 and DDX46 (Prp5) serve similar functions as their yeast counterparts. Here we show that Tat-SF1 is sufficiently conserved that it can supply Cus2 function in the context of yeast splicing. Tat-SF1 binds the Hsh155 ULM *in vitro* (Fig. 1 and 2) via a similar UHM-ULM interface as recently described for human Tat-SF1 and SF3b1 (Loerch et al. 2018). A crystal structure of a complex between Tat-SF1 and Hsh155 further provides detailed inferences about how Cus2 binds Hsh155 (Fig. 3 and 4). Importantly, recombinant Tat-SF1 that lacks a divergent, acidic C-terminal tail can restore the requirement of ATP for pre-spliceosome formation in yeast splicing extracts lacking Cus2 (Fig. 5D). Thus, the conserved RRM/UHM-containing region of Tat-SF1 can fulfill a similar biochemical role as Cus2 in yeast spliceosome assembly suggesting it may play a similar role in mammals.

A striking difference is the affinity these proteins have for each other in different organisms. Whereas SF3b1 ULM5 binds Tat-SF1 with a *K_D_* of approximately 800 nM (Loerch et al. 2018), the ULM of Hsh155 binds Cus2 approximately 25-fold more tightly, and binds Tat-SF1 about three-fold more tightly (Fig. 2). The structure of the UHM of Tat-SF1 and the ULM of Hsh155 (Figs. 3 and 4) suggests several features that could explain the higher binding affinity of the Hsh155-Cus2 interaction compared to SF3b1-Tat-SF1 complexes, including (i) an intermolecular Cus2 D254 – Hsh155 R100 salt bridge as opposed to the intramolecular SF3b1 interaction (e.g. ULM4 E295 – R292), (ii) cation-*π* stacking and hydrogen bond formation between Hsh155 K104 and Cus2 Y252, (iii) additional salt bridges between Cus2 D201/D204 and Hsh155 R95. The difference in affinity may provide different modes of regulation in yeast and humans necessary for accommodating a more divergent set of BP sequences. In humans, the multiple ULMs of SF3b1 interact with a number of UHM containing proteins including alternative splicing factors and the polypyrimidine-tract binding protein U2AF^65^ (Thickman et al. 2006; Corsini et al. 2007; Corsini et al. 2009; Loerch et al. 2014). In fact Tat-SF1 and U2AF^65^ both preferentially bind SF3b1 ULM5 suggesting there may be a sequential series of transient interactions between these proteins during early steps of spliceosome assembly (Thickman et al. 2006; Loerch et al. 2018). Alternatively, the abundance of the U2 snRNPs in these two organisms (5 × 10^5^ U2 snRNPs/cell in Hela cells, 5 × 10^2^ in yeast) may necessitate differences in affinity to maintain proper overall rates of splicing.

The finding that recurrent mutations in SF3b1 during progression of certain cancers alter BP selection in human cells has prompted mechanistic investigations in yeast (Darman et al. 2015; Alsafadi et al. 2016; Tang et al. 2016; Carrocci et al. 2017). The major perturbation detected by transplanting the recurrent cancer mutations into Hsh155 involves interactions between Hsh155 and Prp5. Although the vast majority of the transplanted cancer mutations actually enhance rather than reduce BP fidelity in yeast, other mutations in this region can be selected that reduce fidelity, and these also alter protein-protein interactions between Prp5 and Hsh155. So far, time ordered studies that illuminate the precise steps in spliceosome assembly that are affected have not been done with these SF3b1/Hsh155 mutants, making it difficult to separate specific effects on fidelity from generic effects of loss of function, as described above for Prp5 mutants. If the release of Prp5 serves as the fidelity check of the U2-intron BP helix, then the specific molecular interactions involved may be different, even if the mechanism remains the same.

### How do ATPases exert their function during spliceosome assembly and catalysis?

Genetic studies in budding yeast have been extremely valuable in identifying potential target molecules of DExD/H-box ATPase proteins (e.g. Sub2 release of Msl5-Mud2 from the BP, Prp28 release of U1 snRNA from the 5’SS) (Staley and Guthrie 1999; Chen et al. 2001; Kistler and Guthrie 2001; Hage et al. 2009). However, like most DExD/H-box proteins that function in RNA metabolism, how these proteins exert their functions in RNA splicing is still mysterious. Our work indicates that one outcome of Prp5 ATP binding and hydrolysis is to disrupt binding of Cus2 to Hsh155. However, whether Cus2 release is due to Prp5 directly targeting the Cus2-Hsh155 interface, disrupting Cus2 interaction with U2 snRNA, or unwinding the BSL is still unknown. By binding to the HEAT-repeats of Hsh155 (Tang et al. 2016), Prp5 might be in position to interact with Cus2 and use Hsh155 as a fulcrum to dislodge Cus2 upon ATP-hydrolysis. DExD/H-box ATPases have been implicated in displacing proteins from RNA, and in some cases, independent of RNA duplex unwinding. In vitro, the vaccinia virus DExH ATPase NPH-II can actively remove the U1A protein from a model double-stranded RNA substrate (Jankowsky et al. 2001) and NPH-II and Ded1 were shown to release proteins from model RNP complexes independent of RNA duplex unwinding (Fairman et al. 2004). Alternatively or in conjunction with other factors, Prp5 may weaken Cus2 interactions with RNA and thereby indirectly facilitate its displacement from the Hsh155 complex.

During the transition from commitment complex to the pre-spliceosome, the essential DEAD-box protein Sub2 (human UAP56) is thought to target and release the Msl5-Mud2 heterodimer from the intron branch site (Fleckner et al. 1997; Kistler and Guthrie 2001; Libri et al. 2001; Zhang and Green 2001). However, Sub2 is no longer required for viability when Mud2 is deleted (Kistler and Guthrie 2001), when the Mud2-interacting N-terminus of Msl5 is deleted (Wang et al. 2008), or when the RNA binding surface of Msl5 is mutated (Jacewicz et al. 2015). In humans, these proteins bind through an N-terminal ULM in SF1 (Msl5) and a C-terminal UHM in U2AF^65^ (Mud2) (Selenko et al. 2003). Sequence comparisons with Msl5 and Mud2 suggest they too bind through a ULM-UHM interface consistent with the observation from the Rymond group that deletion of the N-terminus of Msl5 (containing the putative ULM) prevents interaction with Mud2 (Wang et al. 2008). This is analogous to our results presented here with respect to Cus2 and Hsh155 and suggests Sub2 and Prp5 use ATP binding hydrolysis to disrupt consecutive UHM-ULM interactions during early steps of spliceosome assembly. Furthermore, together these data suggest that UHM-ULM interactions may be a previously unrecognized target of DExD/H-box ATPases.

## MATERIALS AND METHODS

### Yeast strains and growth conditions

Yeast strains used and constructed in this study are listed in Table S1 and were grown at 30 °C on YEPD (2% dextrose, 2% peptone, 1% yeast extract) except where noted. Transformations of PCR products and plasmids into yeast cells were carried out using the lithium acetate method according to (Ito et al. 1983). Yeast strains in which the ULM of Hsh155 is mutated (Hsh155^RW-DA^) were constructed by CRISPR/Cas9 mediated editing of the endogenous *HSH155* using a modified approach from that described (DiCarlo et al. 2013) (see below). The *CUS2* ORF was deleted from start to stop codon by transformation and integration of a *natNT2* PCR product with ends homologous to sequences flanking the *CUS2* ORF and conferring resistance to 100 μg/mL nourseothricin (GoldBio) as described (Janke et al. 2004). Strains expressing C-terminal Lea1-TAP were generated by PCR of sequences encoding the TAP tag and the *kanMX4* selectable marker, transformation, and selection on plates supplemented with 200 ug/mL G418 (Sigma-Aldrich) as described in (Janke et al. 2004) and (Rigaut et al. 1999).

### CRISPR/Cas9 editing of *HSH155*

CRISPR/Cas9 mediated editing of the endogenous *HSH155* locus was performed as described in (DiCarlo et al. 2013) with modifications described in (Talkish et al. 2019). To edit the *HSH155* locus in strains BY4741 and Hsh155-13Myc (Pauling et al. 2000), oligonucleotides Hsh155_gRNA_top and Hsh155_gRNA_bot (Table S2) encoding a guide RNA targeting *HSH155* were first annealed and then ligated into the BaeI-digested plasmid p416-TEF1p-Cas9-NLS-crRNA-BaeI. A rescue fragment encoding the R100D and W101A was generated by annealing oligonucleotides Hsh155_RWDA_top and Hsh155_RWDA_bot. The plasmid p416-TEF1p-Cas9-NLS-crRNA-BaeI-Hsh155 and the RWDA rescue fragment were co-transformed into yeast and transformants were selected on SCD-Ura media. After growth on SCD-Ura media, cells that had lost the p416-TEF1p-Cas9-NLS-crRNA-BaeI-Hsh155 plasmid were selected for on media containing 5-fluororotic acid (5-FOA). DNA was extracted from candidate edited clones, PCR amplified using primers that flank the target site, and sequenced at the U.C. Berkeley sequencing center.

### Yeast two-hybrid

Two-hybrid tests between Cus2 or Tat-SF1 and Hsh155 were performed essentially as described in (Yan et al. 1998). Oligonucleotides Gb-155Sph and Gb-AD-155 (Table S2) were used to generate a PCR fragment encoding *HSH155* for Gibson assembly (NEB) into the pACT plasmid. Mutations in *CUS2* and *HSH155* were introduced by site-directed mutagenic PCR using oligonucleotides listed in Table S2. pAS2 plasmids encoding full-length *CUS2*, the *CUS2* UHM (146-285), or *TAT-SF1* (1-383) fused to the *GAL4* DNA binding domain and pACT plasmids encoding *HSH155* fused to the *GAL4* activation domain were transformed into yeast strain y190 and selected for on SCD agar media lacking leucine and tryptophan. Interactions were tested for activation of the *GALUAS-HIS3* reporter by growing cells on SCD agar media lacking leucine, tryptophan and histidine and supplemented with 25 mM 3-amino-1,2,4-triazole (3-AT). Interaction between Tat-SF1 and Hsh155 was tested in a strain in which the *CUS2* open reading frame was deleted. Empty vectors serve as negative controls for reporter activation.

### Cellular extract preparation, SDS-PAGE, and western blot analysis

Cellular lysates from strains expressing Hsh155-13Myc, Hsh155^RW-DA^-13Myc, and Msl5-13Myc were prepared as described in (Ausubel et al. 1994). Samples were mixed with 4x loading dye (250 mM Tris pH 6.8, 40% glycerol, 4% SDS, 4% βME), resolved by SDS-PAGE on 10% acrylamide gels, and transferred to nitrocellulose membranes. Membranes were blocked in 5% milk in TBST (20 mM Tris pH 8, 150 mM Nacl, 0.1% Tween 20) and then incubated overnight at 4 °C with primary antibody diluted in 5% milk in TBST. Mouse monoclonal anti-Myc (9E10) antibody (Sigma-Aldrich) was used at a dilution of 1:5000. Affinity purified rabbit polyclonal anti-Nap1 antibody (a kind gift from Doug Kellogg) was used at a dilution of 1:2000. Primary antibodies were detected using a Li-Cor Odyssey scanner after incubation with IRDye 800CW Donkey anti-rabbit IgG and IRDye 680RD Donkey anti-mouse IgG (Li-Cor) secondary antibodies following the manufacturer’s instructions.

### Splicing extract preparation and *in vitro* splicing reactions

^32^P-radiolabled actin and RP51A pre-mRNA substrates were transcribed *in vitro* using the MEGAscript T7 transcription kit (Invitrogen). Actin pre-mRNA was used in splicing reactions assayed by immunoprecipitation, whereas RP51A pre-mRNA was used in splicing reactions analyzed by native gel electrophoresis. *S. cerevisiae* splicing extracts were prepared using the liquid nitrogen method essentially as described in (Stevens and Abelson 2002) except cells were disrupted using a Retsch MM301 ball mill for 3 minutes at 10 Hz for 5 cycles. ATP depletion and standard splicing reactions were carried out for 20 minutes at 23 °C as described in (Ares 2015) with either 0.4 nM RP51A pre-mRNA or 4 nM actin pre-mRNA. After extracts were depleted of ATP, radiolabeled pre-mRNA and either water or ATP (20 mM final concentration) was added to the reaction and splicing was allowed to proceed for 20 minutes at 23 °C. For reconstitution experiments, recombinant protein was added to the reactions during ATP depletion. Unless otherwise indicated, recombinant Cus2 proteins were added to the reactions to a final concentration of 500 nM and recombinant Tat-SF1 proteins were added to the reactions to a final concentration of (1 µM). To visualize the precursor and product RNAs of splicing reactions, the reactions were diluted in 200 µL of RNA extraction buffer (0.3M NaOAc, 0.2% SDS, 1mM EDTA, 10 ug/mL proteinase K) and incubated at 65 °C for 10 minutes. RNA was then extracted from the reactions with 200 µL of acid phenol, ethanol precipitated, resolved by electrophoresis on 6% acrylamide/8M urea gels, and detected by phosphorimaging. To visualize splicing complexes, reactions were mixed with 2X native loading dye (20 mM Tris/glycine, 25% glycerol, 0.1% bromophenol blue, and 1 mg/mL heparin) and loaded directly on 2.1% agarose gels as described in (Effenberger et al. 2013) and visualized by phosphorimaging. A minimum of two replicates were performed for all in vitro splicing reactions.

### Immunoprecipitations

Splicing complexes were immunoprecipitated using Lea1-TAP from standard splicing reactions containing 40 μl of extract. Dynabeads M270 epoxy beads (Invitrogen) were conjugated to rabbit IgG (Sigma-Aldrich) following the manufacturers recommended procedure. Following splicing conditions, the reactions were diluted in NET-150 buffer (50 mM Tris-Cl pH 7.4, 0.05% Nonidet P-40, 150 mM NaCl) to 400 µL and incubated with 20 µL of IgG-conjugated Dynabeads with gentle rotation for 1 hour at 4 °C. 5% of the reaction was reserved as an input control. After binding, the beads were washed 3 times with ice-cold NET-150 buffer and suspended in 200 µL of RNA extraction buffer (0.3M NaOAc, 0.2% SDS, 1mM EDTA, 10 ug/mL proteinase K) and incubated at 65 °C for 10 minutes. Immunoprecipitated RNA was eluted from the beads and extracted from each sample with 200 µL of acid phenol, ethanol precipitated, resolved on 6% acrylamide/8M urea gels, and visualized by phosphorimaging. The amount of pre-mRNA immunoprecipitated by Lea1-TAP in the absence of ATP was quantified using ImageQuant software (Molecular Dynamics). The mean fold increase and standard deviations of the mean of immunoprecipitated pre-mRNA relative to a wild-type extract was calculated from triplicate experiments from Hsh155^RW-DA^ extracts or extracts lacking Cus2 (Fig. 5B and C). For reconstitution experiments, the mean fold increase of immunoprecipitated pre-mRNA from duplicate experiments was normalized to an extract lacking endogenous Cus2 but containing 500 nM recombinant Cus2 (Fig. 5D-F).

### Recombinant protein expression and purification

*S. cerevisiae* Cus2 for ITC was expressed with a C-terminal His-tag as described (Yan et al. 1998). Cus2 D204K was expressed and purified the same as wild-type Cus2, but was done at the University of California Berkeley QB3 MacroLab. The Cus2 UHM (residues 177-269 of NCBI Refseq NP_014113), human Tat-SF1 (residues 1-360 of NCBI Refseq NP_055315) or the Tat-SF1 UHM (residues 260-353) were expressed in *E. coli* as glutathione-*S*-transferase (GST) fusions using a pGEX-6p variant modified to include a TEV cleavage site for the GST tag. For GST pull-down assays, the Cus2 protein was expressed with an N-terminal His-tag and a GB1-solubility tag. The GST-Hsh155 fragment included residues 86 – 129 (NCBI Refseq NP_014015) since we found that longer constructs of this region were susceptible to proteolysis during purification.

The His-tagged proteins were purified by Ni^+2^-affinity and the GST-tagged proteins were purified by glutathione affinity chromatography. The Tat-SF1 proteins were purified further by heparin-affinity chromatography, whereas the Cus2 UHM was further purified by ion exchange chromatography. For crystallization experiments, the Tat-SF1 UHM was separated from aggregates by size exclusion chromatography in 50 mM NaCl, 15 mM HEPES pH 7.4, 0.2 mM tris (2-carboxyethyl)phosphine (TCEP). For ITC, a synthetic peptide corresponding to the Hsh155 ULM (residues 95 – 109) was purchased at 98% purity (Biomatik Corp).

### GST pull-down assays

The purified GST-fusion proteins (100 μg) were immobilized on glutathione agarose (200 μL, Pierce^™^ Cat. No. 16103) equilibrated with binding buffer (100 mM NaCl, 25 mM HEPES pH 7.4, 10% glycerol, 0.1% NP-40 and protease inhibitors). The GB1-Cus2 protein (500 μg) was added in binding buffer (500 μL) and incubated with the immobilized GST-fusion protein for 30 min. The resin was washed six times with binding buffer. The retained proteins were eluted in SDS-PAGE loading buffer containing 10 mM glutathione and analyzed by SDS-PAGE and Coomassie^®^-Brilliant Blue staining.

### Isothermal titration calorimetry

Protein samples for ITC were dialyzed into 50 mM NaCl, 25 mM HEPES pH 7.4, 0.2 mM TCEP. The peptides were diluted into >200-fold greater volume of the dialysis buffer. The thermodynamics of Cus2 (full-length) or Tat-SF1 (residues 1-360) titrated with Hsh155 ULM peptides were measured using a VP-ITC and fit using Origin 7.0 (MicroCal, LLC). The isotherms were corrected for the heats of dilution using an average of three data points at the saturated plateau of the titration. The apparent stoichiometries of binding were consistent with the formation of 1:1 complexes.

### Crystallization and structure determination

The Tat-SF1 UHM – Hsh155 ULM complex was prepared in a 1:1.2 molar ratio at 18 mg/mL in 50 mM NaCl, 15 mM HEPES pH 7.4, 0.2 mM TCEP and crystallized by vapor diffusion from the JCSG+^™^ screen (Molecular Dimensions) at 277 K. Crystals obtained in the presence of 10% PEG 8K, 8% ethyleneglycol, 100 mM HEPES pH 7.5, were sequentially transferred to 20% glycerol in synthetic mother liquor prior to flash-cooling in liquid nitrogen. Data were collected remotely at Stanford Synchrotron Radiation Lightsource Beamlines 12-2 (Soltis et al. 2008), and processed using the SSRL AUTOXDS script (A. Gonzalez and Y. Tsai) implementation of XDS and CCP4 packages (Table 1) (Kabsch 2010; Winn et al. 2011). The structure was determined by molecular replacement using PDB ID 6N3E with the program Phaser (TFZ-score 11, LLG 2,833) (McCoy et al. 2007), rebuilt in Coot (Emsley and Cowtan 2004), and refined using Phenix (Adams et al. 2010).

## ACKNOWLEDGMENTS

We thank Dr. Sarah Loerch for initiating our collaboration on this project. We also thank Britney Martinez who made initial observations that mutations in the ULM of Hsh155 result in ATP-independent pre-spliceosome formation. We are grateful to Rhonda Perriman, Jen Quick-Cleveland, Stephanie Nystrom, and Logan Mulroney for thoughtful critiques about this work and this manuscript. We appreciate Aaron Hoskins for sharing results prior to publication and for insightful conversations. Thank you to Melissa Jurica for informative feedback and for technical support performing native agarose gel electrophoresis. Thank you to David McPheeters for the Hsh155-13Myc and Msl5-13Myc strains and to Doug Kellogg for generously providing the anti-Nap1 antibody. This work was supported by grant GM040478 from NIGMS to M.A. and grants GM117005 and GM070503 to C.L.K.

---

**Supplemental Figure S1 (to Figure 5A):** Mutation of the ULM of Hsh155 does not affect protein expression or yeast viability. **(A)** Whole cell extracts prepared from cells expressing wild-type or RW-DA mutant Hsh155-13Myc were resolved by SDS-PAGE and assayed by western blotting. An extract containing Msl5-13Myc serves as a control for the specificity of the mouse anti-Myc antibody. Nap1 serves as a loading control. **(B)** Wild-type and Hsh155^RW-DA^ yeast strains grown at 18 °C (5 days), 30 °C, and 37 °C (2 days) on YEPD medium.

**Supplemental Figure S2 (to Figure 5B-D): (A,C,E)** 5% of the *in vitro* splicing reactions immunoprecipitated in Figures 5B-D. **(B,D,E)** Native gel analysis of spliceosome assembly for reactions described in Figure 5B-D. Pre-RP51A was used as a substrate at a final concentration of 0.4 nM. Spliceosomes (sp), pre-spliceosomes (psp), and commitment complexes (cc) are indicated. **(G)** Co-immunoprecipitation of ATP-independent splicing complex with Lea1-TAP in cus2*Δ* extracts reconstituted with wild-type and UHM-mutant Cus2 and Tat-SF1 recombinant proteins. The final concentration of recombinant protein in the splicing reaction is indicated in nM. Tat-SF1 “prep 1” and “prep 2” indicate two independent purifications of recombinant Tat-SF1.

**Table S1:**
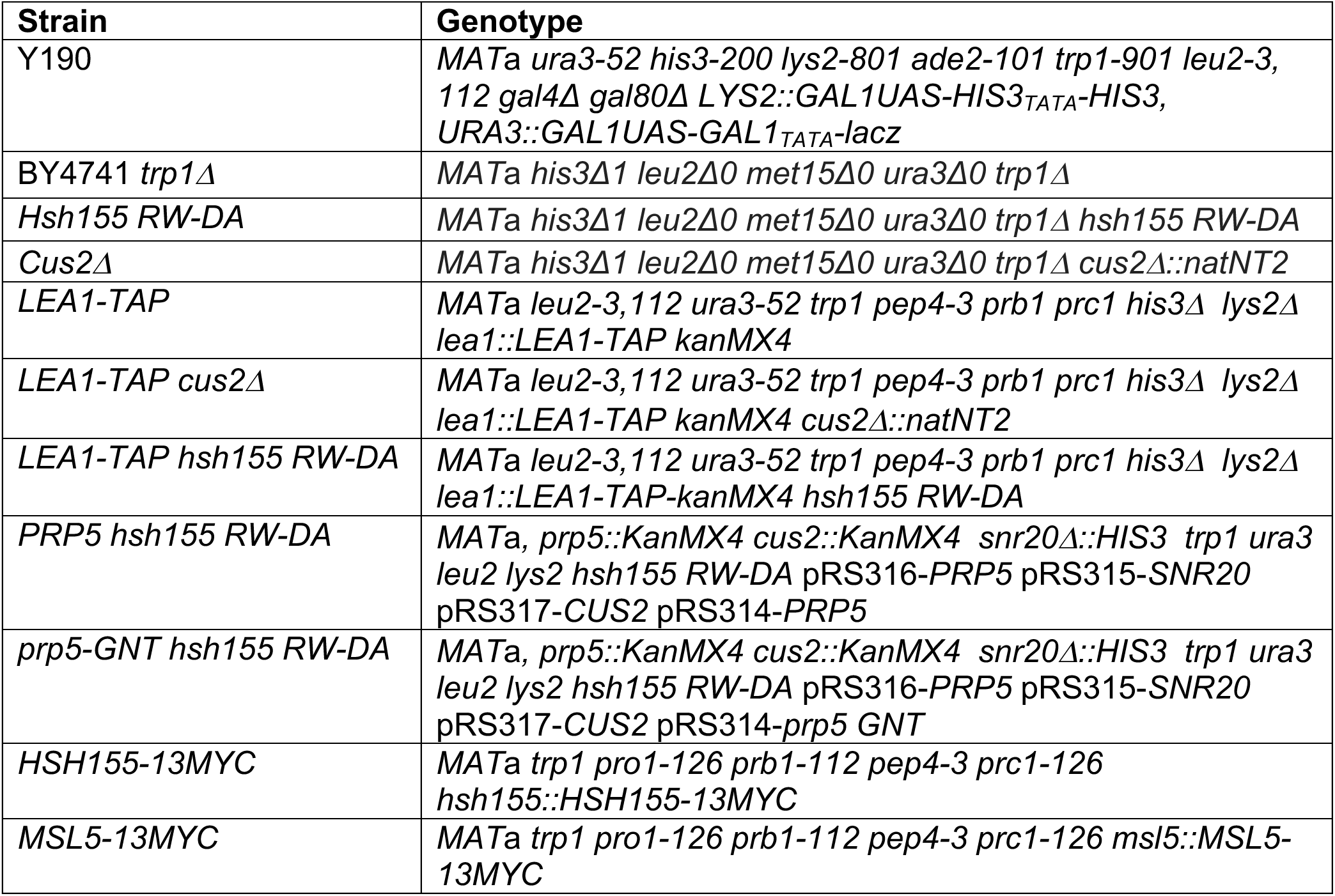
Yeast strains used in this study.

| **Strain** | **Genotype** |
| --- | --- |
| Y190 | *MATa ura3-52 his3-200 lys2-801 ade2-101 trp1-901 leu2-3, 112 gal4Δ gal80Δ LYS2::GAL1UAS-HIS3<sub>TATA</sub>-HIS3, URA3::GAL1UAS-GAL1<sub>TATA</sub>-lacZ* |
| BY4741 *trp1Δ* | *MATa his3Δ1 leu2Δ0 met15Δ0 ura3Δ0 trp1Δ* |
| *Hsh155 RW-DA* | *MATa his3Δ1 leu2Δ0 met15Δ0 ura3Δ0 trp1Δ hsh155 RW-DA* |
| *Cus2Δ* | *MATa his3Δ1 leu2Δ0 met15Δ0 ura3Δ0 trp1Δ cus2Δ::natNT2* |
| *LEA1-TAP* | *MATa leu2-3,112 ura3-52 trp1 pep4-3 prb1 prc1 his3Δ lys2Δ lea1::LEA1-TAP kanMX4* |
| *LEA1-TAP cus2Δ* | *MATa leu2-3,112 ura3-52 trp1 pep4-3 prb1 prc1 his3Δ lys2Δ lea1::LEA1-TAP kanMX4 cus2Δ::natNT2* |
| *LEA1-TAP hsh155 RW-DA* | *MATa leu2-3,112 ura3-52 trp1 pep4-3 prb1 prc1 his3Δ lys2Δ lea1::LEA1-TAP-kanMX4 hsh155 RW-DA* |
| *PRP5 hsh155 RW-DA* | *MATa, prp5::KanMX4 cus2::KanMX4 snr20Δ::HIS3 trp1 ura3 leu2 lys2 hsh155 RW-DA pRS316-PRP5 pRS315-SNR20 pRS317-CUS2 pRS314-PRP5* |
| *prp5-GNT hsh155 RW-DA* | *MATa, prp5::KanMX4 cus2::KanMX4 snr20Δ::HIS3 trp1 ura3 leu2 lys2 hsh155 RW-DA pRS316-PRP5 pRS315-SNR20 pRS317-CUS2 pRS314-prp5 GNT* |
| *HSH155-13MYC* | *MATa trp1 pro1-126 prb1-112 pep4-3 prc1-126 hsh155::HSH155-13MYC* |
| *MSL5-13MYC* | *MATa trp1 pro1-126 prb1-112 pep4-3 prc1-126 msl5::MSL5-13MYC* |

**Table S2:**
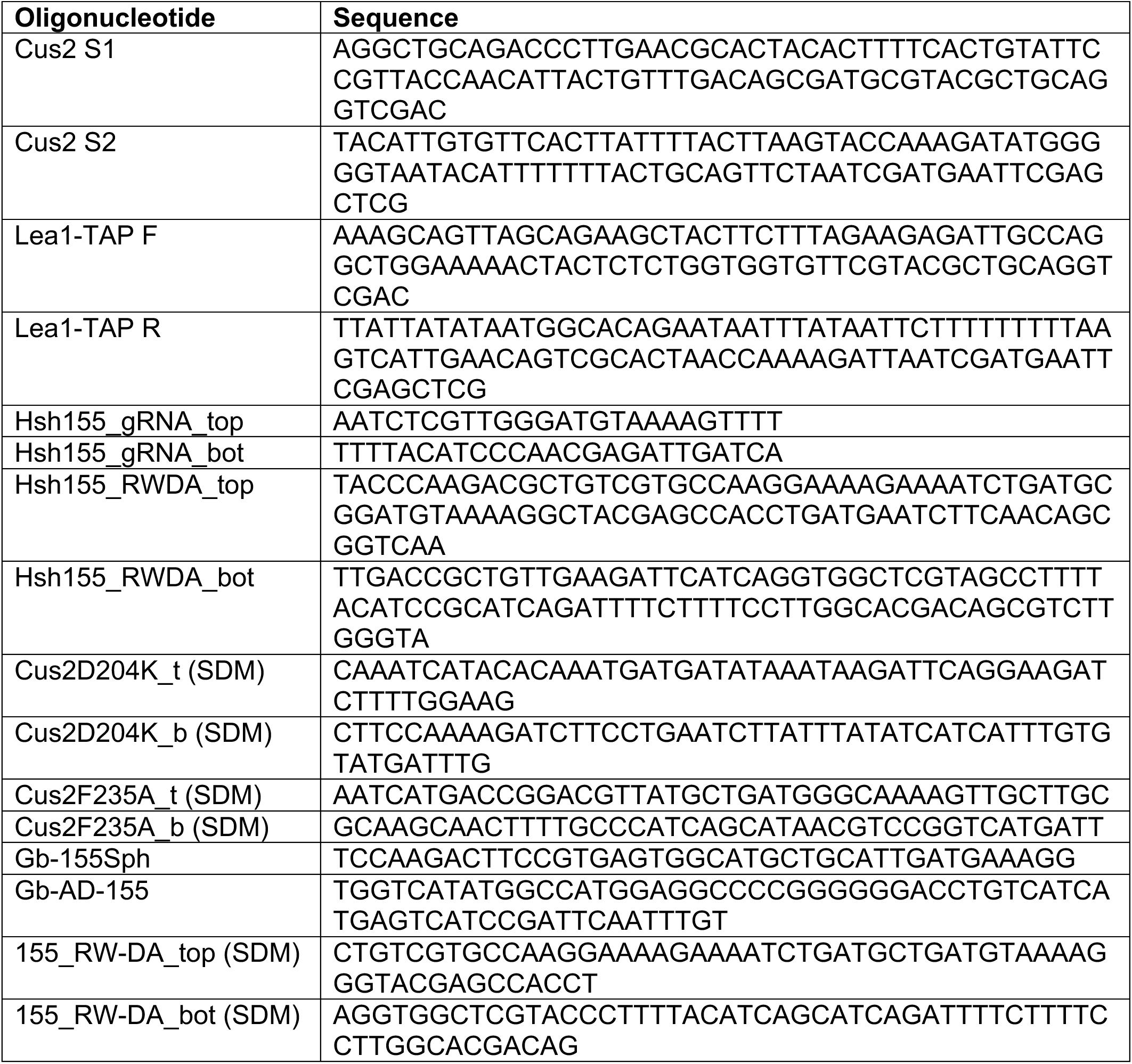
Oligonucleotides used in this study.

