## Supplementary figures and images for "Cus2 enforces the first ATP-dependent step of splicing by binding to yeast SF3b1 through a UHM-ULM interaction"

### Supplemental Figure S1

A

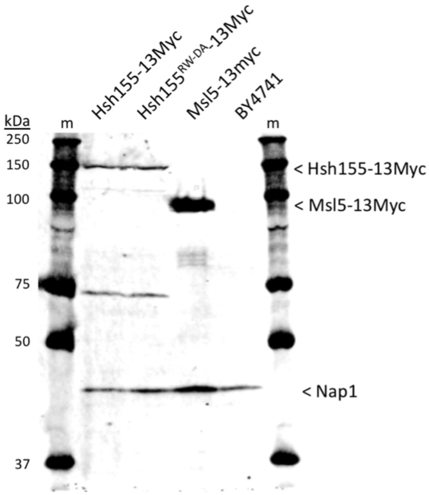


B

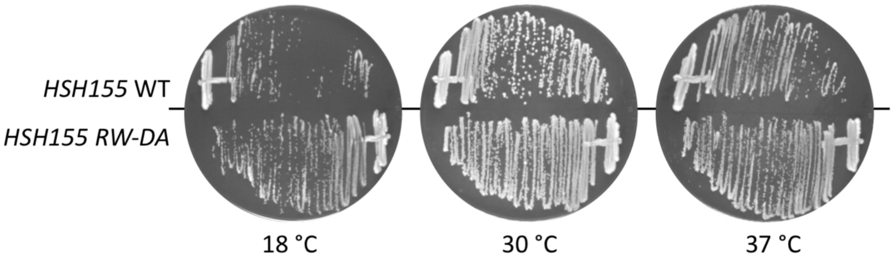


#### Supplementary Figure S1

### Supplemental Figure S2

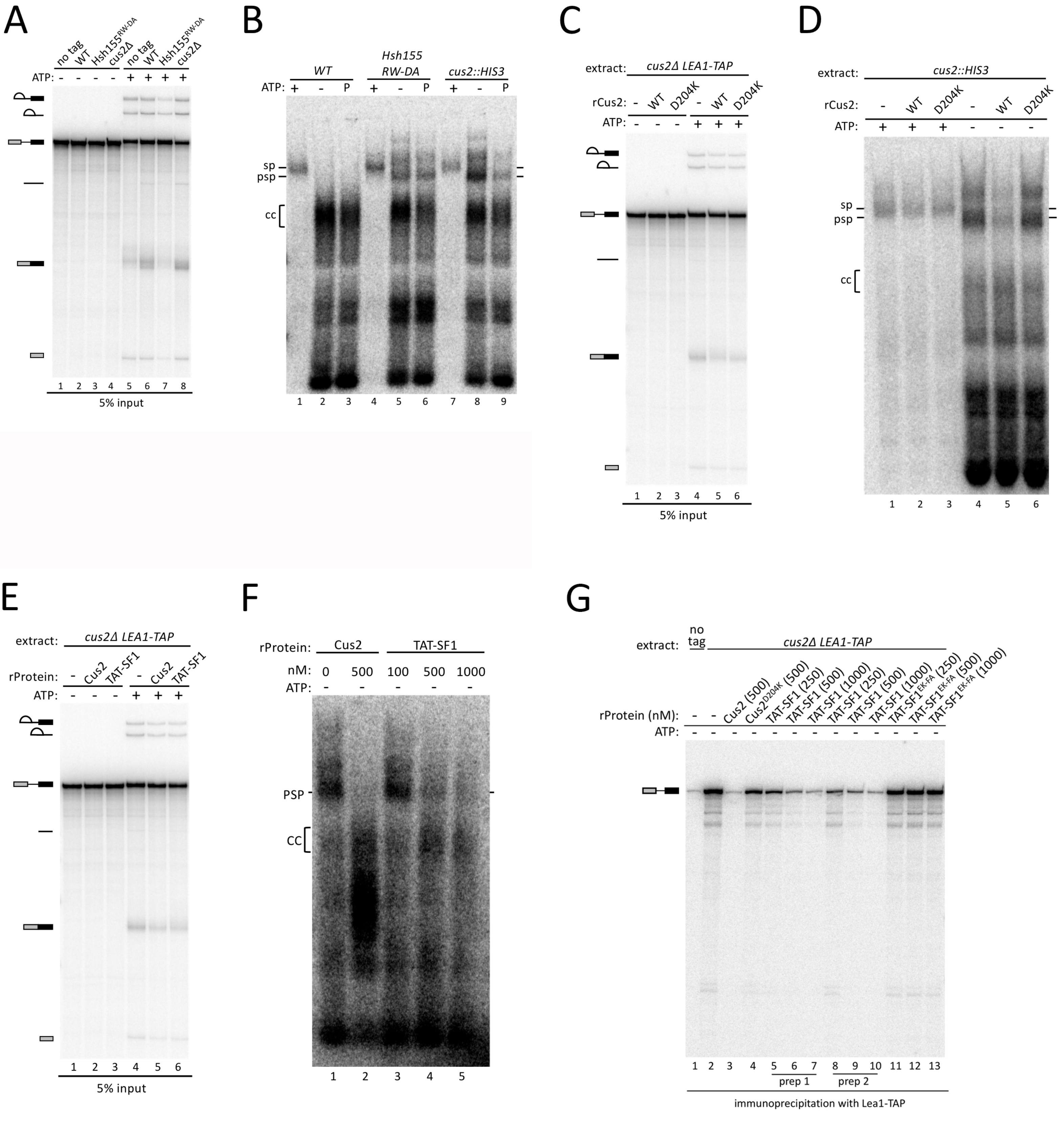


Supplementary Figure S2
